# Cell-cell signaling elicits local Ca^2+^ transients in melanocyte dendrites and dendritic spine-like structures

**DOI:** 10.1101/390542

**Authors:** Rachel L. Belote, Sanford M. Simon

**Affiliations:** Laboratory of Cellular Biophysics, The Rockefeller University 1230 York Avenue, NY, NY 10065

## Abstract

Compartmentalized cytoplasmic fluctuations of Ca^2+^ within dendrites and dendritic spines regulate a variety of neuronal functions. Like some neurons and glia, melanocytes are neural crest derived and possess dendrites (Adameyko et al., 2009; Erickson et al., 1992; Fitzpatrick and Szabo, 1959). Here, we show that primary human melanocytes, when observed *in situ* have extensive dendritic branches with dendritic spines similar to neurons. When co-cultured with primary human keratinocytes, they have local Ca^2+^ transients within these spines and within the dendrites. These are elicited by secreted factors from adjacent keratinocytes. Thus other cell types with dendrites are capable of compartmentalized Ca^2+^ fluctuations in response to cell-cell communication. Furthermore, our observations within intact human skin suggest a more complex communication network between adjacent melanocytes and keratinocytes, and thus a more complex physiology to skin than previous appreciated.

## Introduction

The nervous systems is able to conduct spatially restricted cell-cell signaling via highly specialized dendritic processes that have functional domains and compartments that allow for local response and processing of cell-cell signaling events (Di Castro et al., 2011; Panatier et al., 2011). Chemical synaptic transmission is elicited by an increase of compartmentalized cytoplasmic Ca^2+^ in the presynaptic terminal and often accompanied by a localized increase of cytoplasmic Ca^2+^ within the post-synaptic dendritic spine and/or dendrite (Katz and Miledi, 1965; Llinas et al., 1972; Llinas and Hess, 1976; Llinas et al., 1968; Simon and Llinás, 1985; Spencer and Kandel, 1961; Yuste and Denk, 1995). Cytoplasmic fluctuations of Ca^2+^ within dendrites and dendritic spines regulate a variety of neuronal functions including local synaptic signaling, plasticity, protein translation as well as distal events such as gene transcription(Higley and Sabatini, 2008; Nimchinsky et al., 2002). Subsequently, it has been shown that astrocytes can also respond to cell-cell signalling transmitters with transient compartmentalized increases of Ca^2+^ within their dendritic processes (Di Castro et al., 2011; Panatier et al., 2011). Thus, this type of localized cell-cell communication has been viewed a hallmark of the nervous system. However, dendritic morphologies are not unique to the nervous system. Melanocytes, the pigment producing cells of the skin, share a common lineage with some neurons and glia and also possess dendrites (Adameyko et al., 2009; Erickson et al., 1992; Fitzpatrick and Szabo, 1959). Yet, little is known if melanocyte dendrites share other common features with neuronal dendrites and/or glia processes.

Epidermal melanocytes reside within the basal layer of the epidermis in a ratio of ~1:10 with basal keratinocytes. As the pigment producing cells of the skin, this distribution has led to the concept of the pigmentary unit in which one melanocyte, through its dendrites, has the potential to interact with approximately 30-40 neighboring keratinocytes, the epithelial cells that comprise the bulk of the epidermis. This concept was influenced by studies of in-situ cell density and the observed link between production of melanin in melanocytes and its transfer to neighboring keratinocytes (Fitzpatrick and Breathnach, 1963; Jimbow et al., 1976). Keratinocytes are the most abundant cell type in the epidermis and much work has been done to understand how keratinocytes regulate melanocyte behaviors such as cell proliferation and the production and transfer of pigment throughout the skin (Gordon et al., 1989; Hirobe, 2014). However, much remains unknown about how individual melanocytes interact with keratinocytes and how these two cell types communicate at the single cell level. Given their common dendritic morphology and developmental origin with neurons and glia, we asked if melanocyte dendrites had other structural and functional features of neurons such as a compartmentalized response to cell-cell signaling events.

Examination of epidermal melanocyte morphology in-situ and in a co-culture system revealed a morphological structure on melanocyte dendrites within intact human skin that resembles neuronal dendritic spines. The melanocytes, upon signaling from neighboring keratinocytes, generate locally compartmentalized Ca^2+^ transients within these discrete dendritic spine-like structures as well as within the dendrites themselves. We identified two keratinocyte secreted factors, endothelin and acetylcholine, and the corresponding melanocyte receptors, that are responsible for the localized Ca^2+^ transients within melanocyte dendrites. Our findings that melanocyte dendrites have localized responses to cell-cell signaling events implies a more complex communication network within human skin than previous thought.

## Results

### *In-situ* melanocyte dendrite and dendritic spine morphology

Melanocyte dendrites extend throughout the layers of the epidermis, allowing each melanocyte, both at its soma and dendrites, to interact with multiple neighboring cells. To visualize melanocyte morphology and cell-cell interaction within intact tissue, we used immunofluorescence against the melanocyte specific cell surface receptor KIT proto-oncogene receptor tyrosine kinase (c-Kit) and the melanosome protein tyrosinase-related protein 1 (TRP1). An extensive network of melanocyte dendrites was observed across the basal-proximal layers of the epidermis (Figure 1A, Figure 1 Supplement 1). This is consistent with previous reports that melanocytes have the potential to interact with 30-40 neighboring keratinocytes (Fitzpatrick and Breathnach, 1963; Jimbow et al., 1976). Microinjection of Lucifer yellow into melanocytes of intact human skin (Figure 1B) revealed small protrusions extending from the surface of melanocyte dendrites (Figure 1B, inset). These micron length spine-like processes were observed protruding along the length of the dendrites, but were not visible in the immunofluorescence data, possibly due to incomplete spatial coverage of the melanocyte-specific antigens.

**Figure 1:**
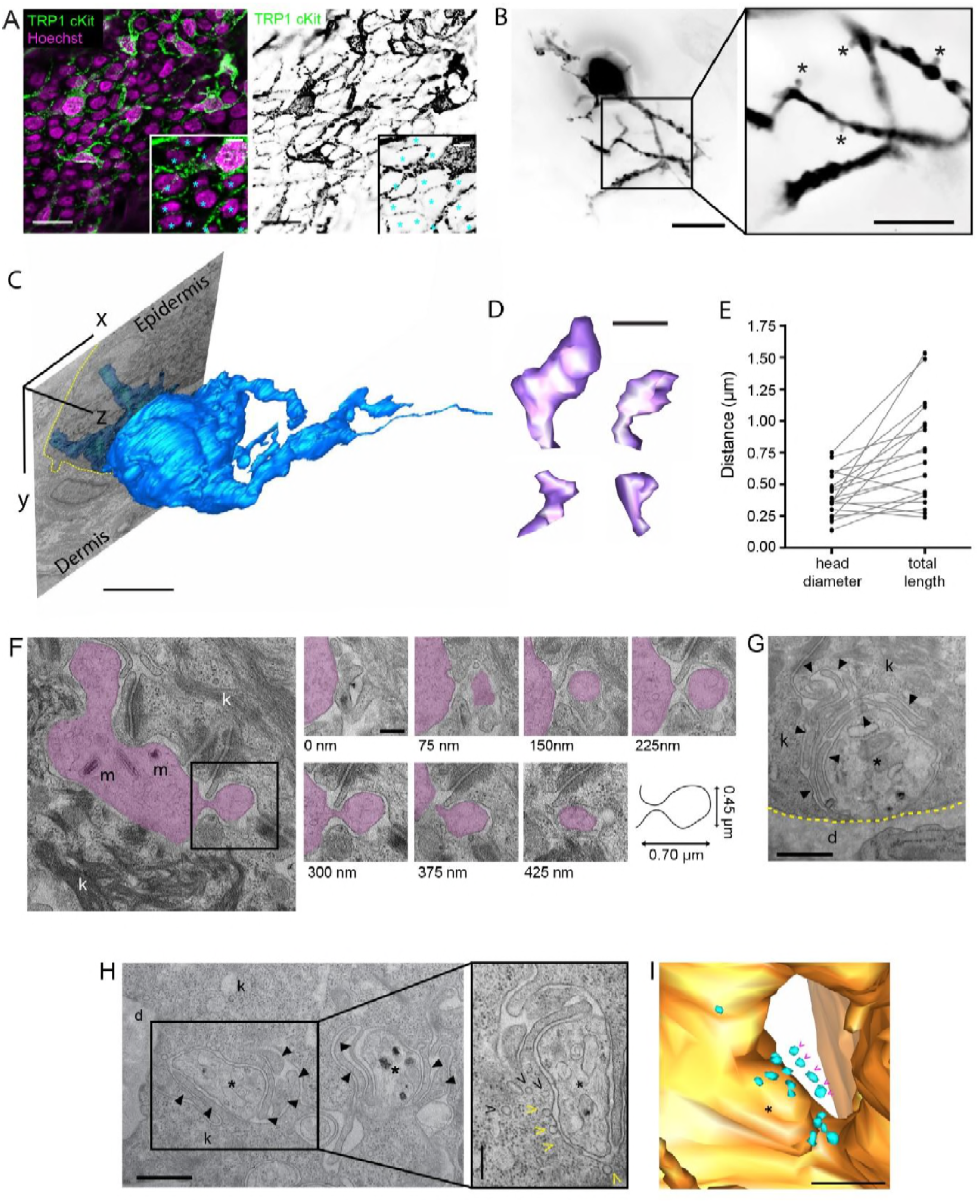
Melanocytes have dendritic spine-like structures. (**A**) Melanocyte dendritic network across the epidermal basal-proximal layer of human neonatal foreskin; non-melanocytic nuclei mark by (*). (**B**), Volume fill morphology of individual melanocyte revealed dendritic spine-like structures (*) on melanocytes in human neonatal foreskin epidermis. (**C**) 3-D reconstruction of melanocyte using 30nm serial FIB-SEM images (**D**), dendritic – spines from FIB-SEM model are similar in shape and (**E**) size to neuronal dendritic spines (n= 19, from 4 cells). (**F**) Melanocyte (purple) dendritic spines are surrounded by adjacent keratinocyte (k) membranes; m, melanosome (**G**) Melanocyte dendrites (*) are surrounded by keratinocyte projections (filled arrow heads); dashed yellow line, basement membrane, d, dermis. (**H** and **I**), Pool of small vesicles (>) within the keratinocyte adjacent to melanocyte dendrite (*). Yellow (>), vesicles fused with the keratinocyte plasma membrane. Scale bars (µm): (**A**), 20; insets, 5 (**B**), 15, inset 10 (**C**), 5 (**D**), 0.5 (**F**), 0.25 (**G**), 0.6 (**H**), 0.6; inset 0.25 (**I**), 0.5.

Using focused ion beam scanning electron microscopy (FIB-SEM) on intact human skin we reconstructed four individual melanocytes amongst 16 surrounding keratinocytes within a volume of 30×27×27 µm at 30 nm z resolution and an XY pixel size of 14-18 nm. An example of one such melanocyte is shown in Figure 1C (See also Figure 1 Supplement 2). Individual melanocytes contacted 7 to 10 keratinocytes within the imaged volume, consistent with the immunofluorescence data. The small spine structures observed protruding from melanocyte dendrites with transmission electron microscopy (TEM) and FIB-SEM (Figure 1D, F and Figure 1 Supplement 3) were indistinguishable from those observed in fluorescence. Thus, the observation of the spines by fluorescence was not an artifact of the preparation and/or microinjection of Lucifer yellow. These structures were similar in shape and size to neuronal dendritic spines *in-vitro* and *in-vivo* (Nimchinsky et al., 2002; Papa et al., 1995; Stuart et al., 2007) (Figure 1E) with head widths of 0.14-0.72 µm and total lengths of 0.24-1.53 µm. The melanocyte spine-like structures (Figure 1F) and the dendrites (Figure 1G,H), were surrounded by and made contact with projections from the keratinocytes similar to the contacts between neuronal dendritic spines and processes from other cells (Ramón y Cajal, 1909). Pools of small vesicles (44–70 nm diameter) were sometimes observed in the keratinocyte where it was juxtaposed to the melanocyte (Figure 1H,I and Figure 1 Supplement 4).

To investigate the melanocyte spine-like structures, we optimized a co-culture system of human primary melanocytes and keratinocytes to mimic physiological conditions of the epidermis (Figure 2A, Figure 2 Supplement 1). We expressed the GAP43-derived fluorescent protein fusion plasma membrane reporters to label melanocytes (eGFP_mem_) and keratinocytes (iRFP_mem_). In this co-culture system, the dendritic arborization of the melanocytes and interactions between the melanocytes and keratinocytes were indistinguishable from those observed *in situ* (Figure 2). Protruding from the melanocyte dendrites were discrete spine-like structures surrounded by projections from adjacent keratinocytes (Figure 2B,D), as observed in intact skin (Figure 1D,F). The spines had head widths from 0.47-1.36 µm and total lengths from 0.55-2.75 µm (Figure 2E). The morphology of some spines were stable, while others exhibited reshaping, moving 0.33-1 µm over an hour (Figure 2B and Figure 2 Supplement 2A,B), similar to dendritic spines in neurons of live mice (Berning et al., 2012). Furthermore, during these movements the melanocyte dendrites remained in contact with the keratinocyte processes indicating a stable interaction (Figure 2 Supplement 2C,D).

**Figure 2:**
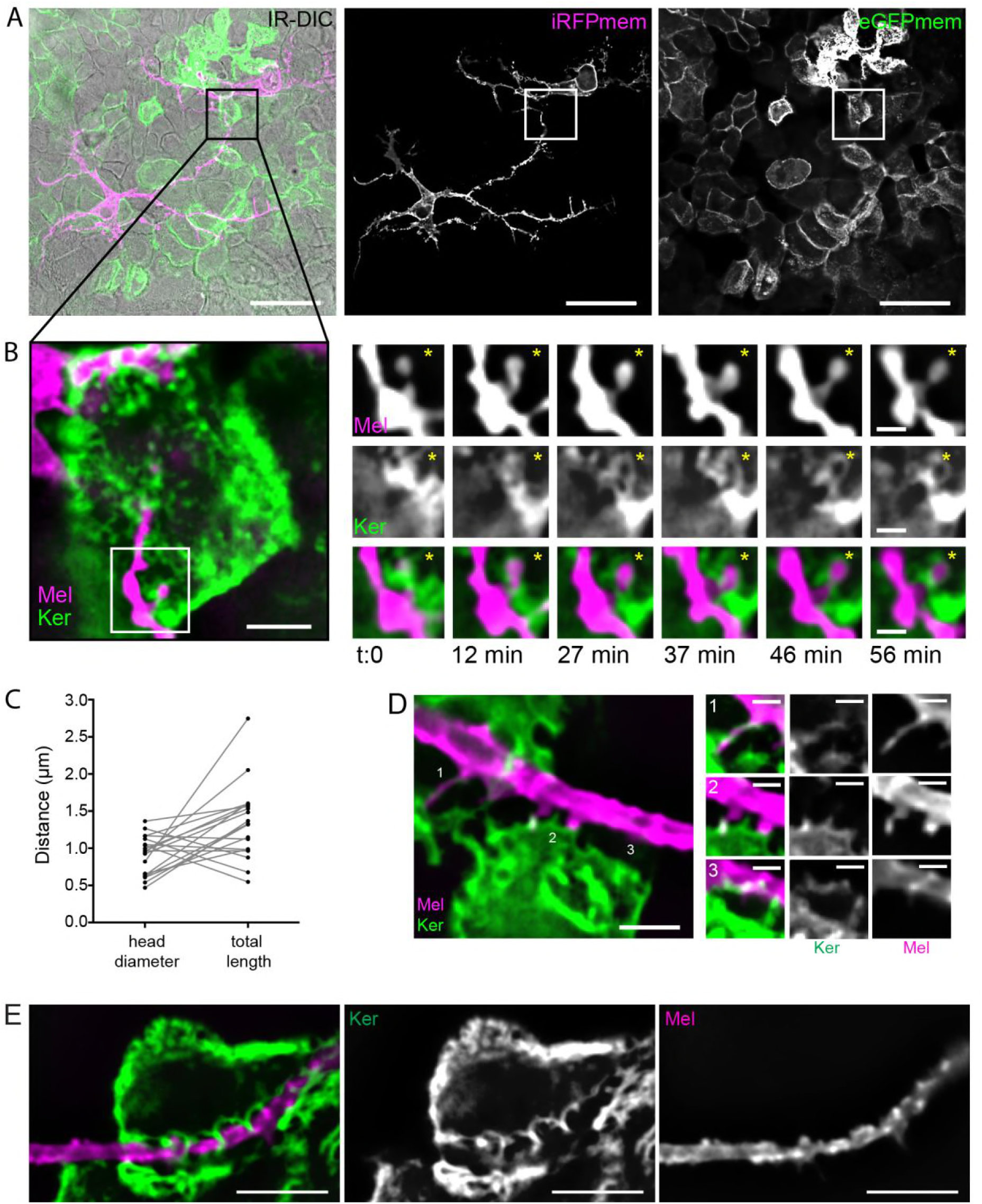
Melanocyte – keratinocyte co-culture recapitulates in-situ cell-cell interactions. (**A**), Melanocytes mimic in-situ morphologies Optimized skin co-culture system is conducive to physiological melanocyte morphologies.(**B**), Some melanocyte (Mel) dendritic spines can have stable morphologies while interacting with adjacent keratinocytes (Ker) while others exhibit dynamic reshaping (**C**), melanocyte dendritic spines in-vitro are similar in size to those in-situ, (**D**), have similar morphologies that are consistent with those observed in neurons and interact with adjacent keratinocyte projections. (**E**), Keratinocyte projections wrap around melanocyte dendrites in the optimized co-culture system. Scale bars (µm): (**A**), 50 (**B**), 5; inset 1.5 (**D**), 5, inset, 2 (**E**),10.

### Dendrites and dendritic spine-like structures compartmentalize Ca^2+^ transients

We tested if the spines on the melanocyte dendrites had a communicative function, similar to neuronal dendritic spines (Higley and Sabatini, 2008; Muller and Connor, 1991; Yuste and Denk, 1995), where synaptic input can elicit local Ca^2+^ influxes (Llinas et al., 1972; Llinas and Hess, 1976; Llinas et al., 1968; Spencer and Kandel, 1961). Using the genetically encoded Ca^2+^ sensor GCaMP6f (Chen et al., 2013), most melanocytes (64±11%, mean ± s.d., n=22 co-culture cultures), when grown in co-culture with keratinocytes, had spontaneous Ca^2+^ transients over a period of 2.5 minutes (Figure 3 and Movie 1). The fraction of melanocytes with calcium transients varied with skin color (59±7% light skin melanocytes and 71±11% medium to dark pigmented skin, Figure 3 Supplement 2A). Of the melanocytes with Ca^2+^ transients, 3±2% had global changes throughout the cell (Figure 3 Supplement 1A-C), while 97±2% exhibited changes localized to sub-regions of the cell (Figure 3A-H, Figure 3 Supplement 1A, D-E). This was irrespective of donor skin color (Figure 3 Supplement 2B).

**Figure 3:**
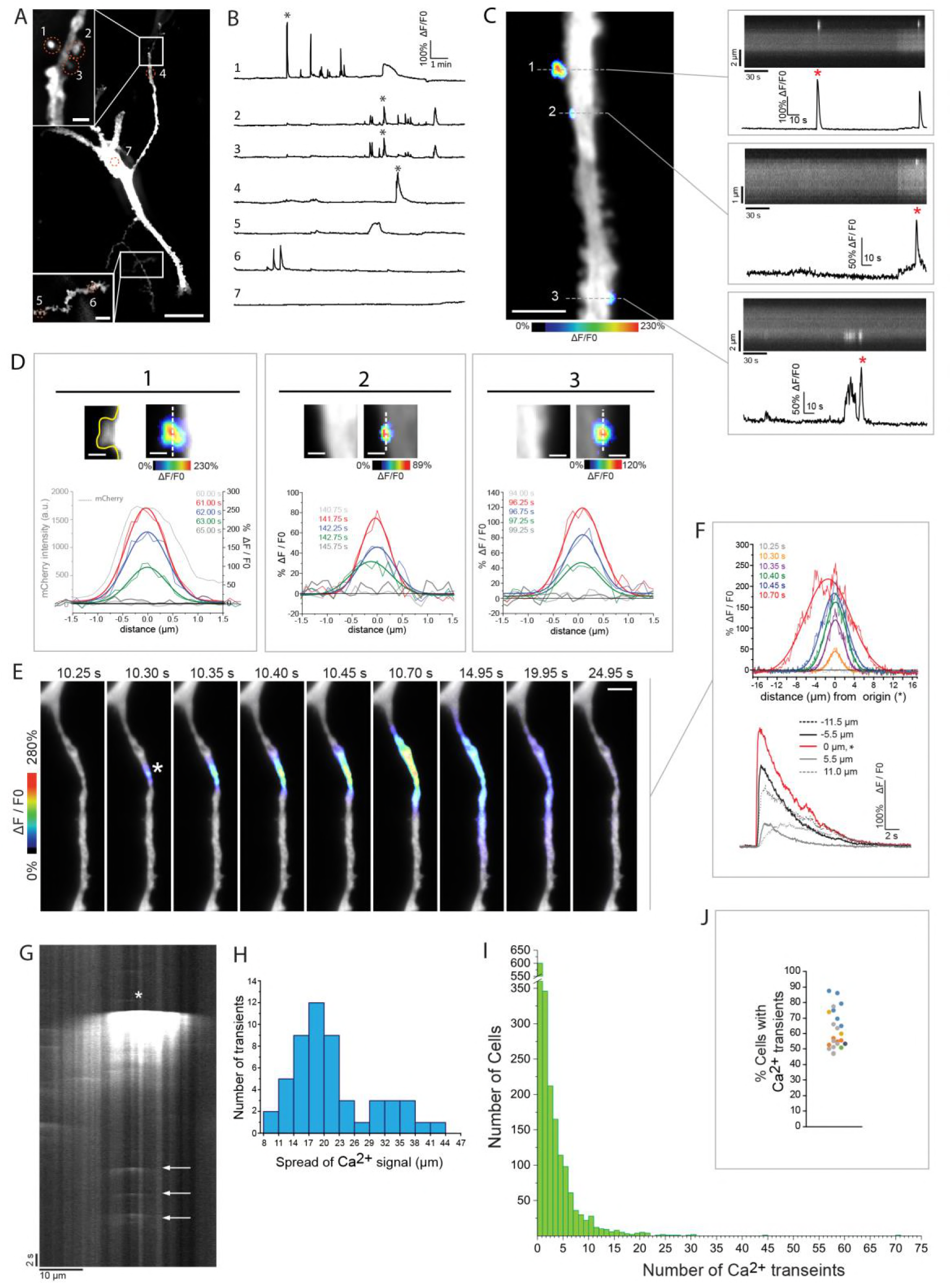
Melanocytes exhibit localized Ca^2+^ transients in the presence of keratinocytes. (**A**), Local Ca^2+^ transients occur at different regions and **(B**), different times within GCaMP6f expressing melanocytes in co-culture with keratinocytes. (**C** and **D**), Ca2+ transients can be restricted in dendritic spines (panel 1) or in confined regions at the dendrite’s surface (panels 2 and 3) (**E** and **F**), Dendritic Ca2+ transients can originate from single detectible site (*) on the dendrite or within multiple regions (**G**), Repetitive Ca^2+^ transients (arrows) at hot spots along the dendrite in **E.** (**H**), Distribution of the spatial spread of the Ca^2+^ signal (FWHM of Gaussian fit) at peak ΔF/F0 intensity for transients within the dendrites (n=53 dendrites from 3 co-cultures from 3 different skin donors) (**I**), Distribution of the number of transients per cell (n=1793 melanocytes from 27 co-cultures from 7 different skin donors). (**J**), The majority of melanocytes have at least one Ca^2+^ transient when co-culture with keratinocytes, (n=22 co-cultures; color indicates skin donor n=6) (FWHM, thick lines). Scale bars (µm): (**A**), 25; top, 2; bottom, 5, (**C**), 5, (**D**), 1 (**E**), 5.

We observed three types of local Ca^2+^ transients within co-cultured melanocytes (Figure 3A-H). The first class was confined within resolvable spine-like structures (Figure 3C and D, panel 1, Movie 2 and 3). The second class occurred within the dendrite (Figure 3C and D panels 2 and 3), which could be the result of an increase in Ca^2+^ within the dendrite or within a spine-like structure not distinguishable by fluorescence. The transients in these first two classes were constrained to a region of <2 µm in diameter (Figure 3C,D). However, the third class affected a larger portion of the dendrite (Figure 3E-H) and resembled, in its extent, the inositol-1,4,5-trisphosphate receptor-mediated dendritic Ca^2+^ transients previously described in pyramidal neurons and Purkinje cells(Finch and Augustine, 1998; Fitzpatrick et al., 2009; Manita and Ross, 2009). Analysis of dendritic Ca^2+^ transients, from 53 cells, across 11 co-cultures from 3 different skin donors showed a range of detectable Ca^2+^ spread from 8-42 µm. The majority (58%) of transients spread over 14-22 µm (Figure 3H), which originated from multiple distinct locations within a region several microns-long of the dendrite (Figure 3 Supplement 1D,E), or from a single point (Figure 3E,F and Movie 4). Additionally, individual dendrites with multiple Ca^2+^ transients over time had hot spots of activity with repeat transients from the same region of the dendrite (Figure 3G). The number of local dendritic transients detected during 2.5 minutes ranged from 0-72 per melanocyte (Figure 3I). The use of a 0.3 N.A., 10× objective limited the analysis to the brightest fluorescence signals and therefore this is likely an under estimate of the frequency.

### Melanocyte dendritic Ca2+ transients are keratinocyte dependent

This high percentage of melanocytes with Ca^2+^ transients (64%±11%) was only observed when melanocytes were co-cultured with keratinocytes. Fewer melanocytes had Ca^2+^ transients when grown with unlabeled melanocytes (16%±2%) or with HEK293T cells (13%±6%) (Figure 4A). The presence or frequency of these calcium transients was not affected by the growth media formulation (Figure 4 Supplement 1A, B). Furthermore, the number of melanocytes with Ca^2+^ transients did not increase when melanocytes were grown in keratinocyte-conditioned media nor when separated from keratinocytes by a semi-porous membrane (Figure 4 Supplement 1C, D). Thus, close proximity to, and possibly direct contact with, keratinocytes was required for the Ca^2+^ transients.

**Figure 4:**
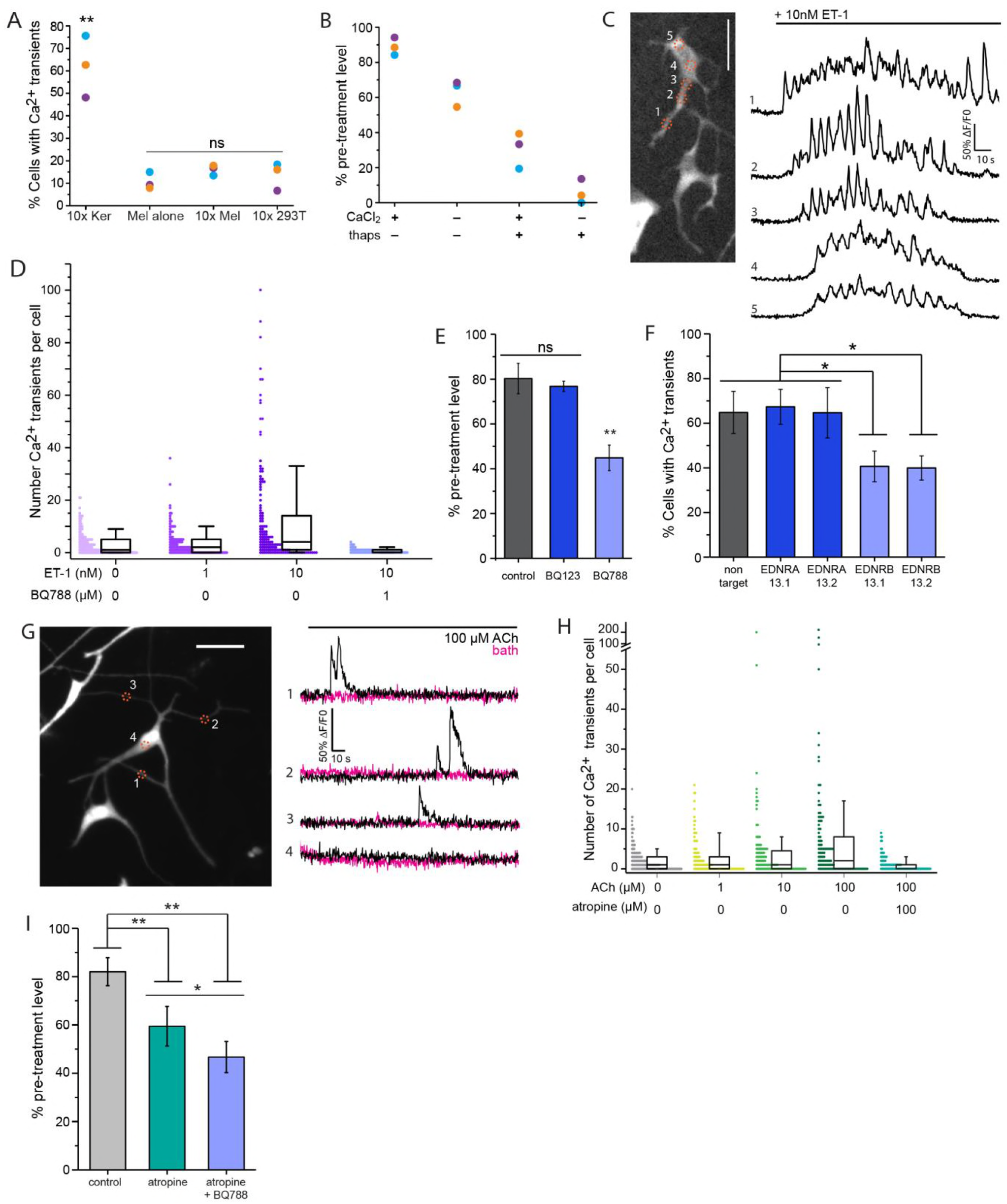
Keratinocyte derived ET-1 and ACh induce Ca^2+^ transients in melanocytes. (**A**), Melanocytes in co-cultures with 10x keratinocytes (10xKer) have significantly more Ca^2+^ transients than those alone (Mel alone), with 10x unlabeled melanocytes (10xMel), or with 10x HEK293T cells (10×293T); n=3 co-cultures per condition; colors indicate different skin donors. One way ANOVA F(3,8)=29.05, p-value=0.00012; post-hoc Tukey means comparison to other culture conditions, ** p-value <0.0005; ns, not significant: p-value > 0.05. (**B**), Both external and internal Ca^2+^ stores contribute to Ca^2+^ transients in co-cultured melanocyte. n=3 cultures per condition. Colors represent different skin donor. Significant by One-way ANOVA with post hoc Tukey means comparison (F(3,8)= 67.8, p-value <0.00001, Tukey: all p-values <0.01). (**C**), Ca^2+^ oscillations in melanocytes co-cultured with keratinocytes in response to 10 nM ET-1 (black line: addition of 10nM ET-1 to imaging bath), ΔF/F0 plots corresponding to numbered regions on the melanocyte. (**D**), ET-1 increases the number of Ca2+ transients per co-cultured melanocyte via the ET_B_ receptor (n=203,141,141,62 cells from 3, 2, 2, 1 different patient derived and matched co-cultures, respectively). Significant by Kruskal-Wallis ANOVA (H= 67.63, 3 d.f., p-value = 1.37 x10^-14^). Box: 25%, 75%; Whiskers: 10 % - 90%. (**E**), ET_B_ (BQ788) and not ET_A_ (BQ123) inhibition significantly reduces the number of melanocytes with Ca^2+^ transients, n = 6, 3, and 3 co-cultures, respectively derived from 6 different donors. One way ANOVA F(2,9)=39.38, p-value=0.000035; post-hoc Tukey means comparison to other culture conditions: ** p-value <0.0005; ns, p-value > 0.05; data represented as mean ± s.d. (**F**), Only knockdown of melanocyte ET_B_ reduced the number of melanocytes with Ca2+ transient in co-cultures. One way ANOVA F(4,10)=8.21, p-value=0.0033; post-hoc Tukey means comparison to other culture conditions, * p-value <0.05; data represented as mean ± s.d., n= 3 different skin donor derived co-cultures. (**G**), 100µM ACh elicits local transient Ca^2+^ response in melanocyte dendrites. Black line: addition of ACh (black ΔF/F0 plot) or imaging media (bath, magenta ΔF/F0 plot). (**H**) ACh, via mAChRs, increases the number Ca^2+^ transients per co-cultured melanocyte (n=243,128,128,115, 131 cells from 4, 2, 2, 2, 2 different patient derived and matched co-cultures, respectively). Significant by Kruskal-Wallis ANOVA (H= 41.32, 4 d.f., p-value = 2.371 x10^-8^). Box: 25%, 75%; Whiskers: 10 % - 90%. (**I**), mAChR antagonist antropine reduces the number of co-cultured melanocytes with Ca^2+^ transients, n = 5, 3, and 3 co-cultures, respectively, derived from 5 different donors. One way ANOVA F(2,8)=28.75, p-value=0.00022; post-hoc Tukey means comparison, ** p-value <0.005; *, p-value = 0.10; data represented as mean ± s.d. Scale bars: 100 µm.

To determine the source of Ca^2+^ responsible for the dendritic transients, we quantified the number of Ca^2+^ transients before and after the removal of external CaCl_2_ and/or addition of 1 µM thapsigargin, which releases calcium from internal stores(Lytton et al., 1991). The percent of cells with Ca^2+^ transients was reduced by either the removal of external CaCl_2_ or addition of thapsigargin (63±4% and 30±5% of pretreatment level, respectively) compared to the control (88±5% of the pretreatment, Figure 4B), with the combination of removal of CaCl_2_ and addition of thapsigargin having the greatest effect (7±4% of the pretreatment level). This was also true for the number of Ca^2+^ transients per cell (Figure 4 Supplement 2).

### ET-1, a keratinocyte derived paracrine factor, contributes to dendritic Ca^2+^ transients

Endothelins, a family of 21 amino acid peptides, play a critical role in melanocyte development, maturation, and homeostasis within the epidermis (Reid et al., 1996). Epidermal keratinocytes produce and secrete endothelin 1 (ET-1)(Yohn et al., 1993) which can elicit increases of Ca^2+^ in the cell bodies of melanocytes (Kang et al., 1998). Addition of 10nM ET-1 to co-cultures produced localized recurring Ca^2+^ transients across the entire melanocyte, including dendrites and the cell body (Figure 4C, Figure 4 Supplement 3A and Movie 5). The percent of melanocytes with a response to ET-1 was dose dependent (Figure 4 Supplement 3B). The number of transients per melanocyte also increased with the addition of 10nM ET-1 (Figure 4D). The ET-1-induced increase of Ca^2+^ in melanocytes co-cultured with keratinocytes was blocked by pre-incubation with a selective inhibitor of endothelin receptor B (ET_B_), BQ788, but not an inhibitor of endothelin receptor A (ET_A_), BQ123 (Figure 4E, Figure 4 Supplement 3C,D and Movie 6). This was also true of melanocytes in mono-culture (Figure 4 Supplement 4).

The localized Ca^2+^ transients in melanocytes elicited by ET-1 (Figure 4C and Movie 5) resembled those seen in co-cultures of melanocytes and keratinocytes without the addition of exogenous ET-1 (Figure 3 and Movie 1). To test if the local Ca^2+^ transients in melanocytes resulted from release of endothelin by keratinocytes, we treated co-cultures with endothelin receptor antagonists. Co-cultures treated with the ET_B_ antagonist had a significant reduction in the percent of melanocytes with Ca^2+^ transients but no change in Ca^2+^ transients in co-cultures treated with the ET_A_ antagonist (Figure 4F). To confirm that the effects were due to inhibition of the ET_B_ on melanocytes and not the consequence of inhibition of ET_B_ on keratinocytes, we used DsiRNA to knock down either ET_A_ or ET_B_ in melanocytes prior to co-culture with keratinocytes. Consistent with the endothelin receptor antagonist data, knock down of ET_B_ but not ET_A_, reduced the number of co-cultured melanocytes with Ca^2+^ transients (Figure 4F).

### Heterogeneous melanocyte response to acetylcholine

Inhibition of ET_B_ did not completely eliminate all Ca^2+^ transients in co-cultured melanocytes. In addition to producing and releasing endothelin, keratinocytes also possess the necessary cholinergic machinery to synthesize, release and degrade acetylcholine (ACh), a classic neurotransmitter(Grando et al., 1993). Furthermore, melanocytes express muscarinic receptors (mAChR) and respond to the mAChR specific agonist, muscarine (Buchli et al., 2001). Addition of ACh increased the number of transients per cell in co-culture (Figure 4G,H and Figure 4 Supplement 5A) and mono-culture (Figure 4 Supplement 6A), with some having over 100 transients in 2.5 min, a 10-fold increase compared to control conditions (Figure 4H and Movie 7). This increase in frequency of Ca^2+^ transients, in both co- and mono-cultured melanocytes, was blocked by atropine, a selective muscarinic ACh receptor antagonist (Figure 4H and Figure 4 Supplement 6B). ACh increased the frequency of calcium transients, in those cells that already showed activity, but had no effect of number of melanocytes with Ca^2+^ transients (up to100µM, Figure 4 Supplement 5B). Atropine decreased the number of co-cultured melanocytes with transients to 54±8% of the pretreatment value (Figure 4I), indicating that endogenous ACh release caused melanocyte Ca^2+^ transients. Addition of BQ788 with atropine, further reduced the number of melanocytes with transients to 47 ± 0.6% of the pretreatment value (Figure 4I).

## Discussion

Neuronal dendritic spines were first characterized over a century ago (Ramón y Cajal, 1909), they were first shown to have regenerative excitatory activity 50 years ago (Spencer and Kandel, 1961), and electrogenic calcium responses 40 years ago (Llinas et al., 1972; Llinas and Hess, 1976; Llinas et al., 1968). However, it is only recently that it has been demonstrated that, by compartmentalizing signaling, dendrites and dendritic spines allow for spatial control of processes such as protein translation, enzymatic activation and recruitment of membrane receptors (Higley and Sabatini, 2008; Nimchinsky et al., 2002). Here we show that another cell type with dendritic processes is capable of compartmentalized signaling. The localized Ca^2+^ transients within melanocyte dendrites and their spines adds a functional parallel between melanocytes and cells of the nervous system which has profound implications for how melanocytes receive and process information from their surroundings. It also suggests that there may be a more complex communication network between melanocytes and keratinocytes. These results show that one melanocyte, through its dendritic arbor, physically interacted with multiple keratinocytes and keratinocytes contacted multiple melanocytes. This suggests that the melanocyte-centered “units” within the epidermis may not be independent.

Furthermore, we found that keratinocytes also have small cytoplasmic projections both *in-situ* and in culture which made contact with melanocytes (Figure 1G,H and Figure 2D,E). Similar structures were first described by Palade in amphibian skin at keratinocyte-keratinocyte junctions and have been referred to as microvilli by other groups when discussing phagocytic events in keratinocytes (Ando et al., 2012; Farquhar and Palade, 1965). However, not much is known about their function or molecular composition and it remains to be determined as to what class of cell extension they belong. The keratinocyte projections we observed wrapped around individual melanocyte dendrites (Figure 1G), made contact with melanocyte dendritic spines (Figure1F, Figure 1 Supplement 3), and could form physical barriers between adjacent dendrites (Figure 1H). This local arrangement is similar to that of glial cells of the nervous system (Auld and Robitaille, 2003; Chong et al., 1999). Thus, keratinocyte projections may serve similar functions such as physically supporting the architecture of the melanocyte network across the skin as well as maintaining local signaling niches. In the work presented here, we showed that keratinocyte secreted endothelin and ACh caused a localized response in melanocyte dendrites (Figure 4). Within the context of a multicellular tissue, the physical envelopment of melanocyte dendrites by the keratinocyte projections could facilitate localized delivery and recycling of secreted factors similar to what occurs between neurons and glia at chemical synapse (Auld and Robitaille, 2003; Chung et al., 2015). Further investigation into the molecular components of the keratinocyte projections as well as the stability and time scale of their interactions with melanocyte dendrites and spines would provide further insight into their function.

### Melanocyte dendritic functional domains

While previous work has demonstrated that melanocytes can have changes in Ca^2+^ in response to keratinocyte-derived paracrine factors, only melanocyte cell body Ca^2+^ concentration was monitored (Imokawa et al., 1997; Kang et al., 1998). Here were show that in co-culture with keratinocytes, melanocyte dendritic Ca^2+^ transients originated from distinct regions within the melanocyte dendrite with subsequent events occurring within the same region as well as at specific dendritic spines (Figure 3). This was also true of ET-1 and ACh induced Ca^2+^ transients (Figure 4 Supplement 3 - 6). These data are indicative of functional domains along melanocyte dendrites in which receptors and linked signaling cascades are restricted to, or active at, specific regions of the dendrite. However, further investigation is required to determine exactly what factors contribute to the observed spatial distribution of localized Ca^2+^ transients, including, but not limited to, Ca^2+^ buffering proteins, ER Ca^2+^ availability, spatial distribution of IP3 receptors, as well as store operated calcium and calcium induced calcium response machinery.

Through its dendritic arbor, one melanocyte physically interacts with multiple keratinocytes. The presence of functional domains across the entire dendrite would allow melanocytes to locally modulate cell processes within a specific region of the dendrite. One possible function for this type of spatial control could be local regulation of the production and transfer of melanin. The observation that, on average, a higher percentage of Ca^2+^ transients were observed in melanocytes from derived from dark skin than from light skin (Figure 3 Supplement 2) suggests they might regulate local melanin production and/or melanosome transfer. This is consistent with previous work that showed an overall increased cytoplasmic Ca^2+^ enhances mono-cultured melanocytes’ response to melanogenic stimuli (Carsberg et al., 1995).

### Melanocyte heterogeneity

Interestingly, not all melanocytes responded to ACh with detectable increases in Ca^2+^ transients, which indicates that at the individual level there is heterogeneity in response to ACh (Figure 4 G,H and Figure 4 Supplement 3). ACh acts through two different classes of cell surface receptors: muscarinic receptors (mAChR), a family of G protein coupled protein receptors, and nicotinic receptors (nAChR), a family of multi-subunit ligand gated ion channels (Hulme et al., 1990). Our data is consistent with previous work that showed melanocytes, at a population level, express mAChRs (Buchli et al., 2001). While it is possible that nicotinic receptors are expressed in melanocytes, the level to which atropine, the mAChR antagonist, blocks the ACh response in melanocytes suggests nAChRs are not the main responders to ACh. It is important to note that cytosolic Ca2+ increase upon AChR stimulation is not ubiquitous and depends on coupled g proteins, in the case of mAChRs (Hulme et al., 1990). Therefore, the observed heterogeneity could be the result of a) heterogenous AChR expression among melanocytes and/or b) variability of receptor subtype or g coupled proteins within individual melanocytes.

### Local vs global response to stimulus

Compartmentalization of second messengers allow for spatial control of biochemical response to external stimuli that provide the cell a mechanism for local regulation of a variety of processes including protein translation, enzymatic activation and recruitment of membrane receptors (Higley and Sabatini, 2008; Nimchinsky et al., 2002). It is possible that the level of stimulus can determine the difference between a localized response and a global cell response in melanocytes. Given that melanocytes interact with many different cells within the epidermis, it is possible that melanocytes use their dendrites to compartmentalize responses to local stimuli whereas stimulus from multiple adjacent cells would initiate a global response across the whole cell. Keratinocyte secreted endothelin elicited a localized Ca^2+^ transient response in melanocytes (Figure 6E-G) whereas addition of 10nM ET-1 directly to melanocytes both in mono and co-culture lead to Ca^2+^ transients across the entire melanocytes (Figure 6C and Figure S6A,E).

ET-1 modulates, in a dose dependent manner, melanogenesis, mRNA transcription, and proliferation of melanocytes in mono-culture (Imokawa et al., 1997; Imokawa et al., 1995; Tada et al., 1998). The transcription of mRNA and cell proliferation require transduction of signals into the nucleus. On the other hand, melanogenesis can be locally regulated via alterations in the activity of melanin synthesizing enzymes on melanosomes and globally, throughout the whole cell, through changes in the amount of melanosome specific proteins. ET-1 has been shown to both up regulate activity of tyrosinase, a key enzyme in melanin synthesis, as well as increase tyrosinase mRNA transcription in melanocytes (Imokawa et al., 1997). The ability to threshold a response to stimulus, such as local vs global Ca^2+^ increases, offers a possible mechanism by which melanocytes can make decisions regarding such outcomes.

UV radiation has a dose dependent effect on skin pigmentation through activation of intrinsic melanocyte responses as well as up-regulating multiple melanogenic keratinocyte-derived paracrine factors (Bellono et al., 2013; Hirobe, 2005). It is possible that basal levels of skin pigmentation are regulated through local, sub-threshold signaling along melanocyte dendrites, whereas doses of UV radiation that activate a tanning response (increase in melanin production, melanin transfer and transient proliferation of melanocytes) cause an increase in signaling that exceeds this threshold producing a whole cell response.

Neuronal dendritic spines are dynamic structures involved in short- and long-term plasticity at synapses. It is possible that the dendritic spine-like structures on melanocyte dendrites function similarly to their neuronal counterpart: integrating signals to determine the states of melanin production while providing a way to adapt to long-term responses to UV radiation.

---

**Figure 1 supplement 1:**
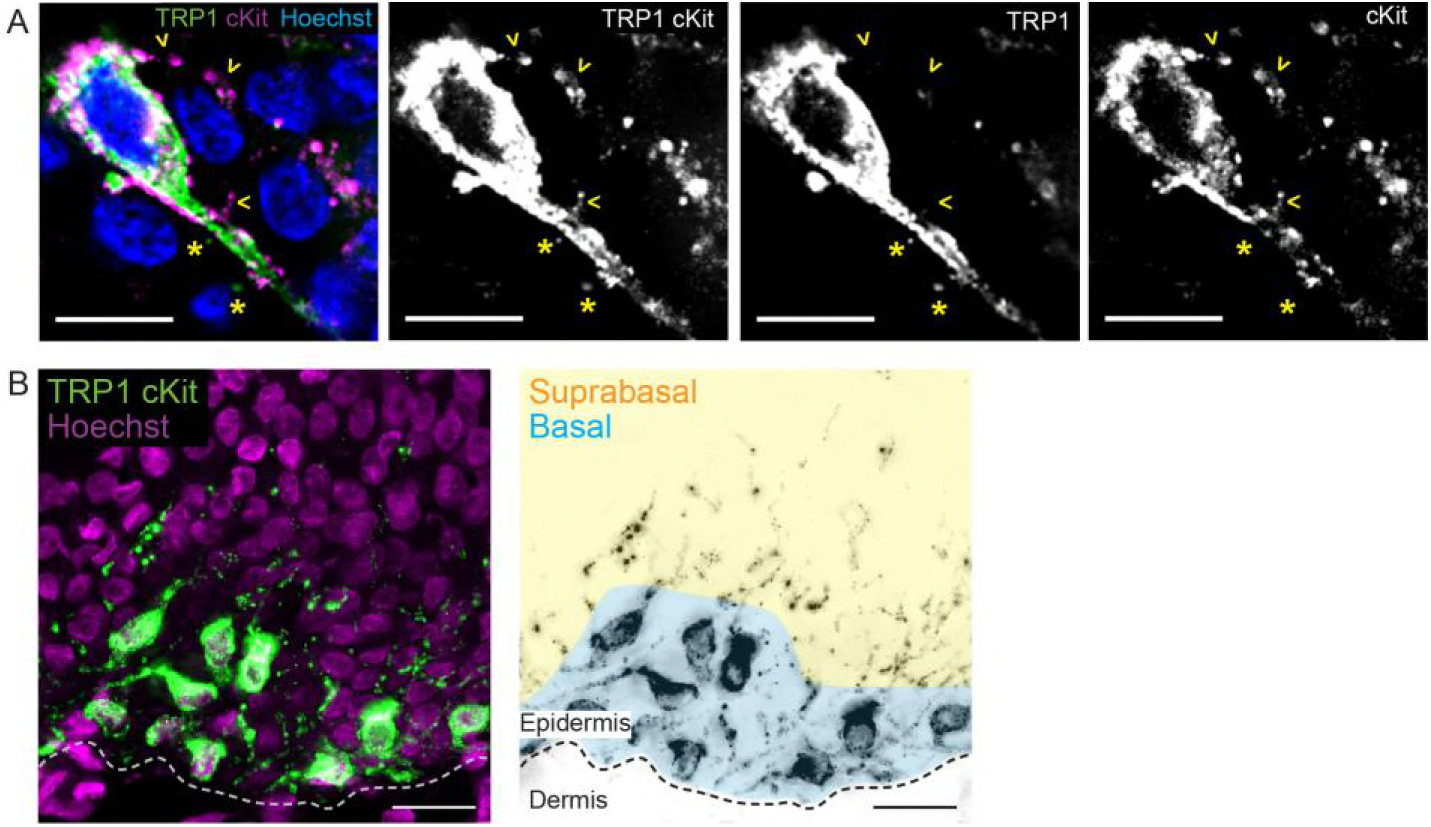
(**A**) Melanocyte-specific antibody cocktail (α-TRP1 and α-c-Kit) provides extensive labeling of melanocytes in human epidermis. Yellow * indicates region only recognized by α-TRP1, yellow > indicates regions only recognized by α-c-Kit. Scale bars, 10 µm. (**B**) Angled cross section of intact neonatal foreskin. Dotted line demarcates the basement membrane. Blue region highlights melanocyte cell bodies in the basal layer. Orange indicates suprabasal layers. Scale bars, 20 µm

**Figure 1 supplement 2:**
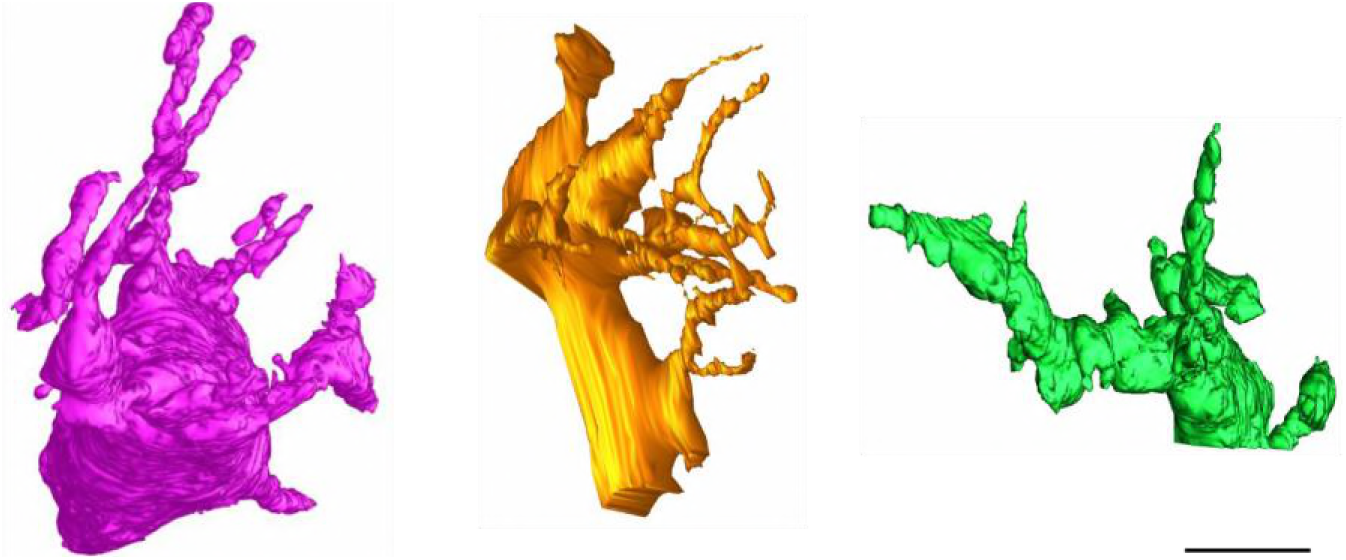
Morphology of the 3 additional individual melanocytes from the FIB-SEM 3-D reconstruction model. Scale Bar, 5µm.

**Figure 1 supplement 3:**
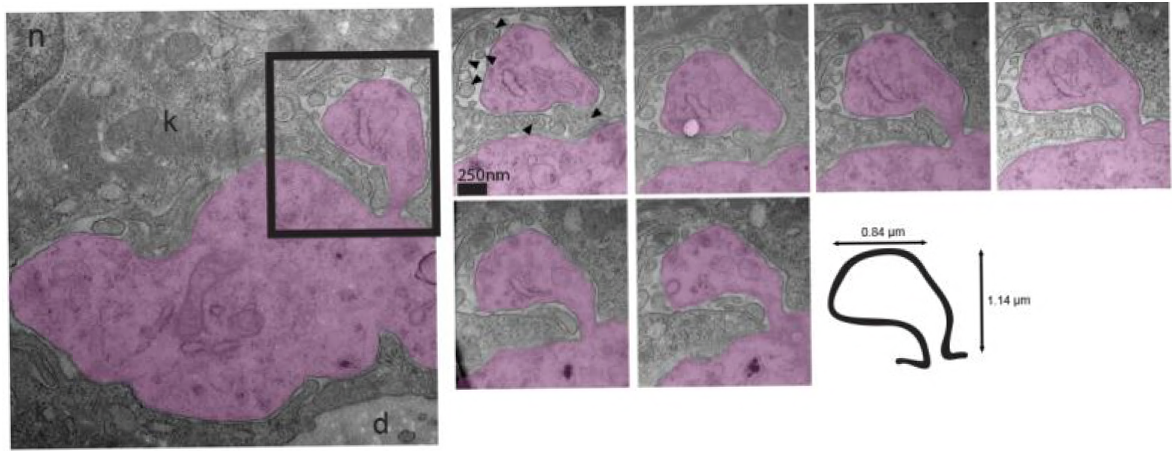
Representative spine-like structure from intact neonatal foreskin with illustration depicting the morphology and size of the spine-like structure. Melanocyte dendrite in purple; n, nucleus; d, dermis; arrow head, keratinocyte processes. TEM serial sections (50-100 nm). Scale bar 250 nm.

**Figure 1 supplement 4:**
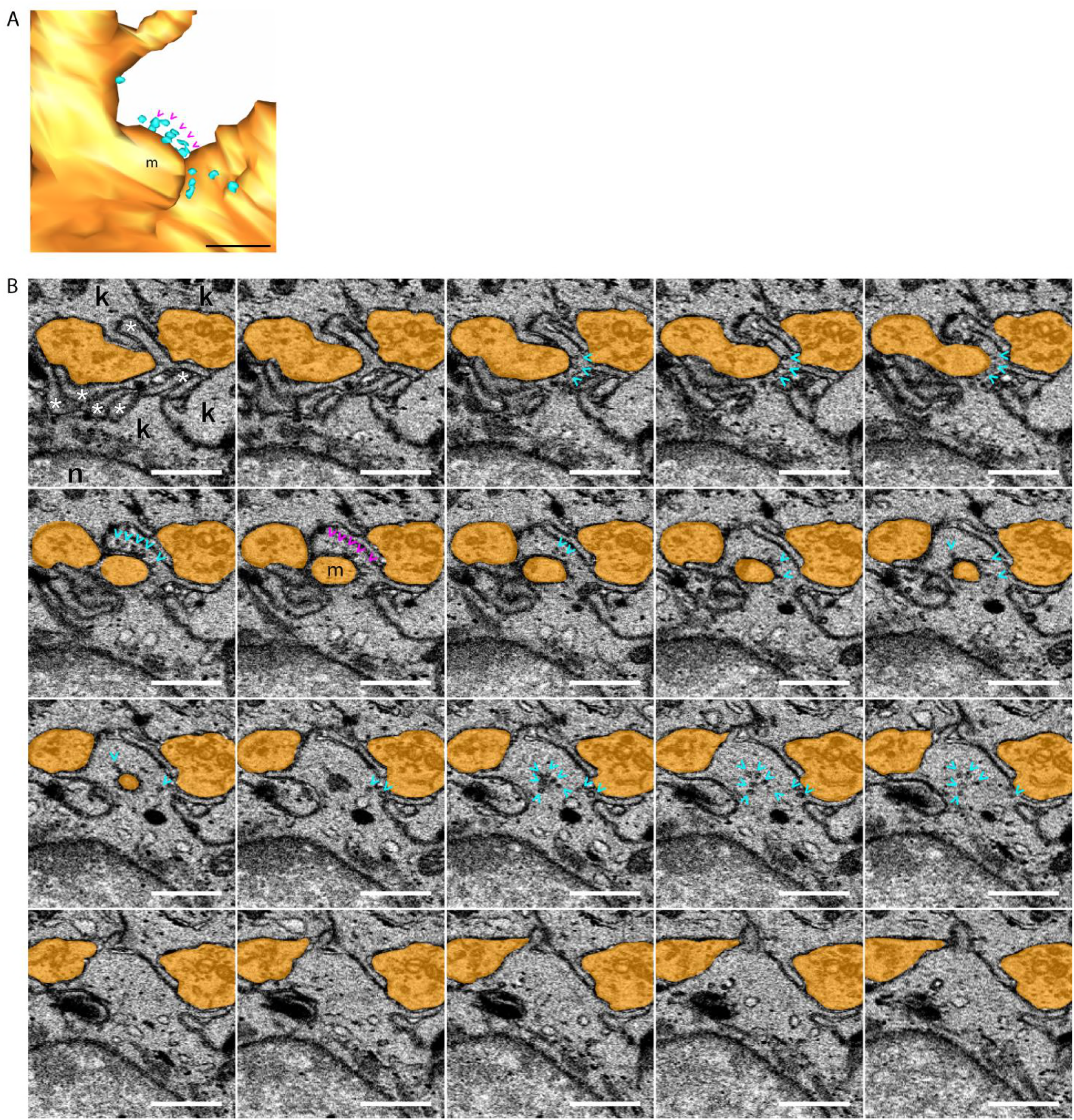
Pooled vesicles in keratinocyte adjacent to melanocyte dendrite. (**A**), 3D reconstruction of melanocyte m1 from figure 1D. magenta (>) corresponds to vesicles in Figure 1I ; m corresponds to region of melanocyte that is adjacent to keratinocyte vesicles in (**A**). Scale bar 0.5 µm. (**B**), serial 30nm z sections from FIB-SEM data that were used to generate the 3D reconstruction in Figure 1. *, keratinocyte projections; k, keratinocyte; n, nucleus; orange: melanocyte m1 ; cyan (>), vesicles. Scale bar 1 µm

**Figure 2 supplement 1:**
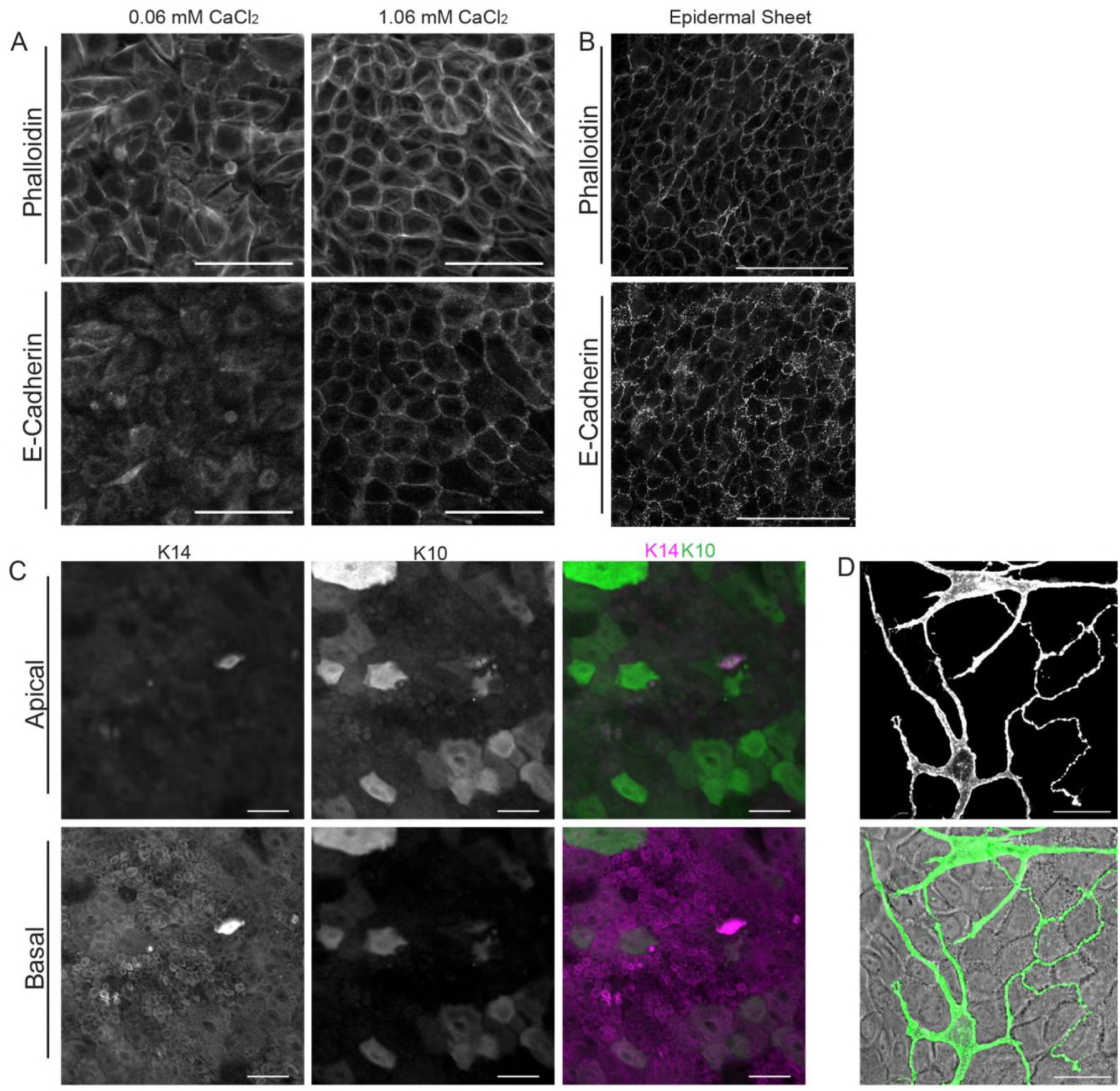
Characterization of melanocyte – keratinocyte co-culture (**A**), Representative images of cytoskeleton morphology, assessed by phalloidin binding of F-actin, and E-cadherin expression in melanocyte–keratinocyte co-culture grown in EpiLife media + HKGS with 0.06 mM CaCl_2_ or 1.06 mM CaCl_2_. Scale bars, 100 µm. **(B**), F-actin cytoskeleton morphology and E-cadherin localization in a fixed freshly isolated neonatal foreskin epidermis. Scale bars, 50 µm. **(C**), Keratin 14 (K14) positive keratinocytes localize to the basal layer of the culture while keratin 10 (K10) positive keratinocytes localize at the apical surface in the melanocyte–keratinocyte co-culture grown in EpiLife media + HKGS for 17 hours in 0.06 mM CaCl_2_ then for 48 hours in 1.06 mM CaCl_2_. Scale bars, 200 µm. **(D**), Melanocytes expressing iRFPmem grown in optimized co-culture system. Scale bar 30 µm.

**Figure 2 supplement 2:**
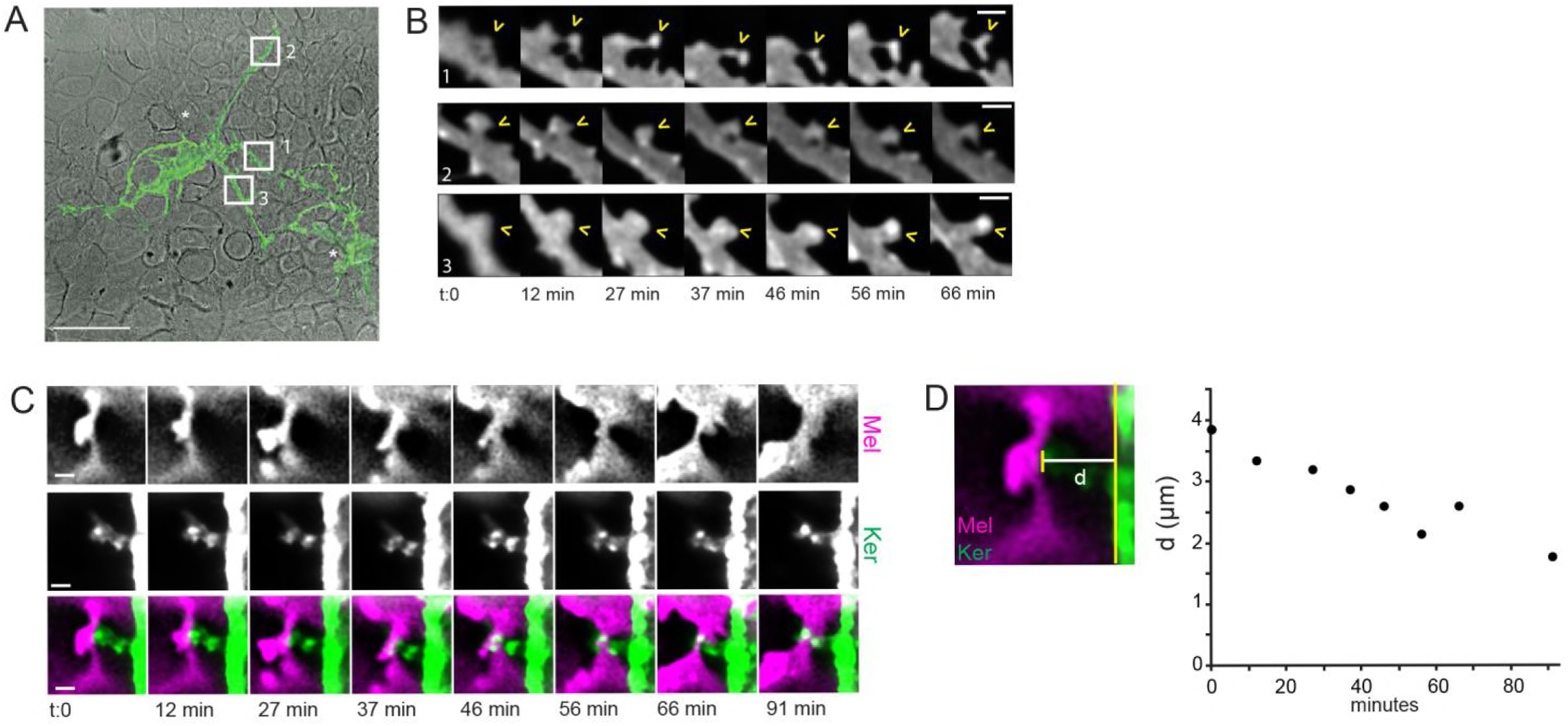
Morphological changes in melanocyte spine-like structures over time. (**A**), iRFPmem labeled melanocytes (green) in confluent co-culture with keratinocytes (IR-DIC), * melanocytes. Scale bar, 50 µm. (**B**) Time-lapse images of spine-like structures, marked by open arrow heads, from the white boxes 1, 2 and 3 in (**A**). Scale bars, 2 µm. (**C**), Keratinocyte projections remain in contact with melanocyte dendrite during changes in morphology and placement. Live cell time-lapse imaging of a keratinocyte projection interacting with a melanocyte dendrite. Scale bars, 2 µm. (**D**), Melanocyte dendrite displacement relative to the keratinocyte cell body during interaction with the keratinocyte projection in (**C**). Distance (d) was measured between the farthest edge of the melanocyte dendrite (short yellow line) to the closest edge of the keratinocyte cell body (long yellow line).

**Figure 3 supplement 1:**
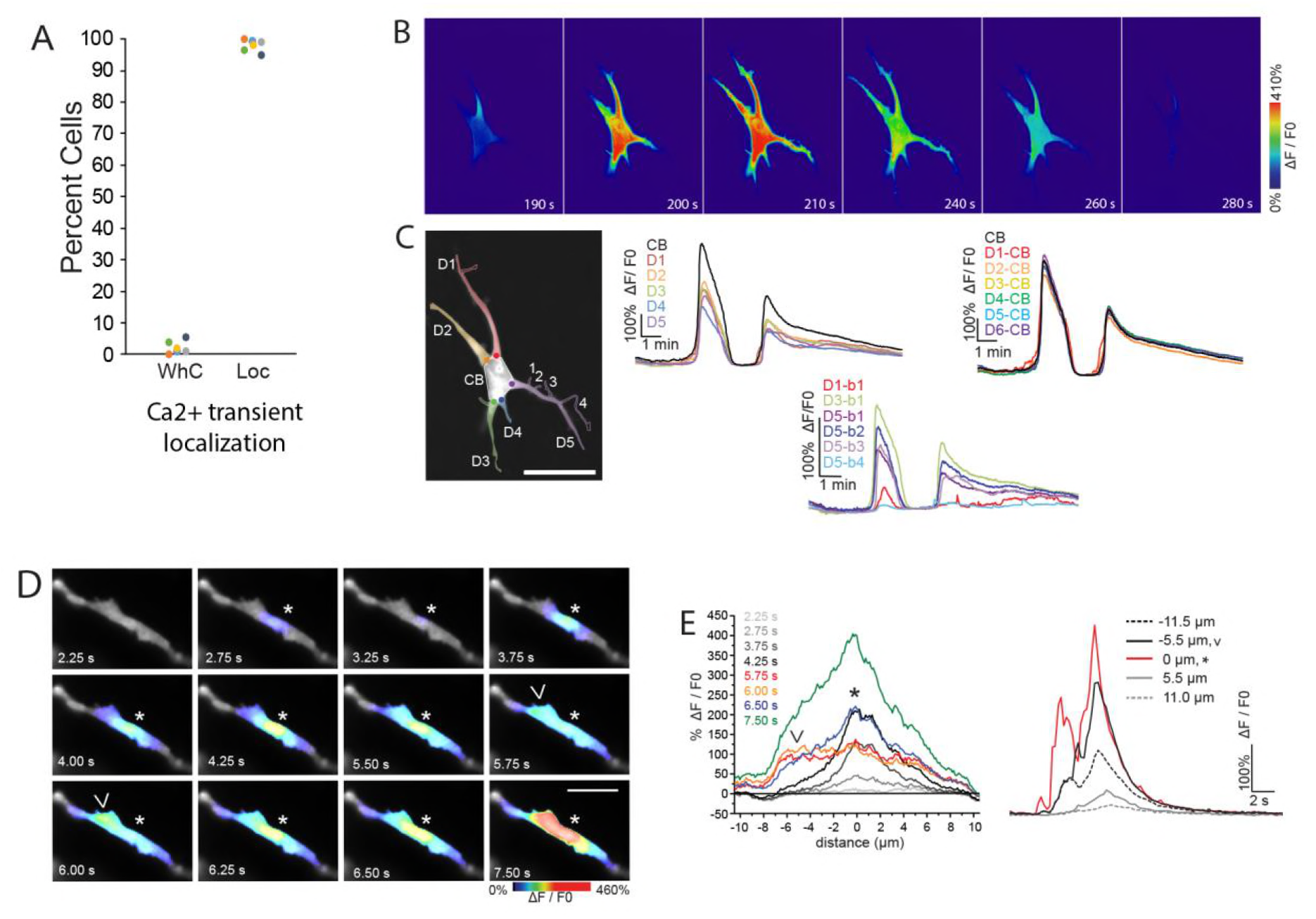
Whole cell and local Ca^2+^ transients in melanocytes. (**A**), Percent melanocytes with global whole cell (WhC) or local (Loc) Ca^2+^ transients. Data from Figure 3J pooled by skin donor. (**B**), Representative images of whole cell Ca^2+^ transient with (**C**), corresponding ΔF/F0 plot from designated regions: CB, cell body; DX, primary dendrites where X indicates dendrite number; DX-CB, dendrite – cell body junction; DX-by, secondary dendrites, y indicates secondary branch number. (**D** and **E**), Spatiotemporal properties of dendritic Ca2+ transients. (**D**), Ca^2+^ transients originating at two adjacent regions * and >. (**E**), Ca^2+^ spread at corresponding times marked in (**D**) and time course of Ca^2+^ transients at indicated distances from the initiation site * where region > is represented at -5.5 µm with solid black line. Scale bars (µm): (**C**), 100, (**D**), 5.

**Figure 3 supplement 2:**
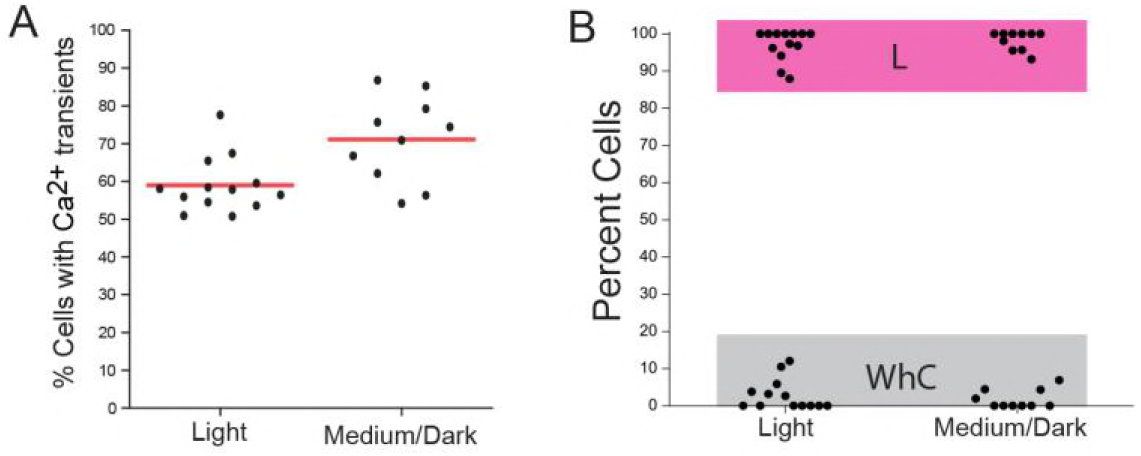
Comparison of Ca^2+^ transient occurrences in co-cultures derived from light skin versus medium to dark skin. (**A**) Percent melanocytes expressing GCaMP6f that have one or more Ca^2+^ transients in co-culture with keratinocytes. Melanocytes and keratinocytes were derived either from light skin or from medium to dark skin. Transients were observed in 58.9% ± 7.4% (mean ± s.d., red line) of light skin melanocytes, and in 71.1% ± 11.3% of medium to dark skin melanocytes. Two Sample T-test p-value < 0.01. (**B**) Percent of melanocytes with whole cell (WhC) Ca^2+^ transients or with local (L) Ca^2+^ transients. (A and B) Data from main text Figure 3 grouped by donor skin color.

**Figure 4 supplement 1:**
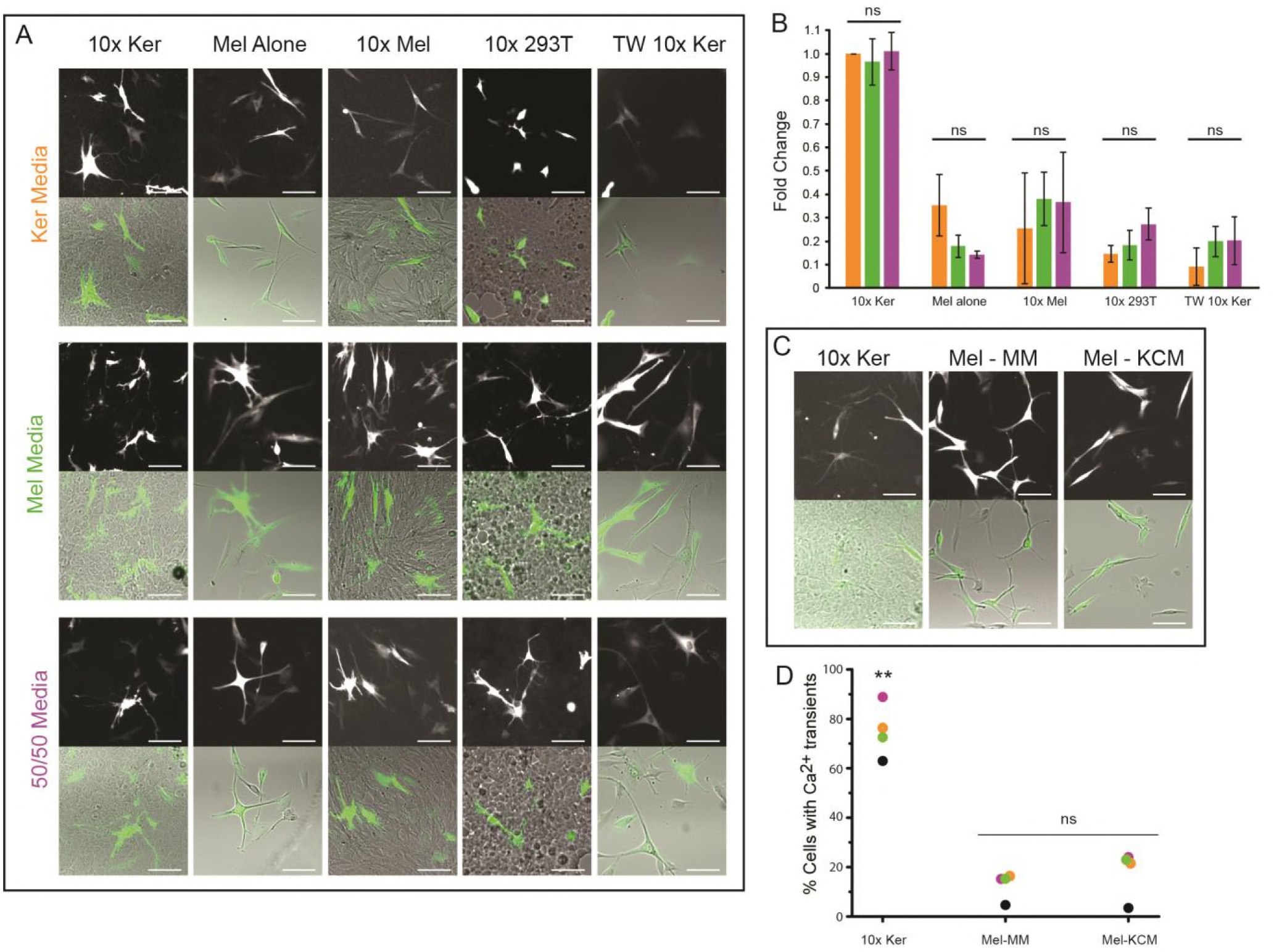
Comparison of Ca^2+^ transient occurrences in co-cultures derived from light skin versus medium to dark skin. (**A**), Representative images of GCaMP6f-expressing melanocytes in co-culture or mono-culture grown in the indicated conditions: 10x Ker: normal co-culture conditions of GCaMP6f melanocytes with 10x keratinocytes; Mel alone: GCaMP6f melanocytes cultured alone; 10x: GCaMP6f melanocytes cultured with 10x unlabeled melanocytes; 10x 293T: GCaMP6f melanocytes cultured with 10x HEK293T cells; TW 10x Ker: GCaMP6f melanocytes cultured with 10x keratinocytes separated by a porous membrane. Ker media: keratinocyte growth media. Mel Media: melanocyte media. 50/50 Media: media containing 1:1 keratinocyte: melanocyte growth media. (**B**), Fold change in the percent of GCaMP6f-expressing melanocytes with Ca2+ transients in the indicated conditions: keratinocyte growth media (orange), melanocyte growth media (green), 50/50 media (purple). One-way ANOVA analysis within each culture type showed no difference between growth media; ns, not significant. Data represented as mean ± s.d.; n ≥ 3 biological replicates per culture and media condition; Fold change is the ratio of the given culture condition to the corresponding biological replicate’s 10x Ker in keratinocyte growth media. (**C**), Representative images of GCaMP6f-expressing melanocytes in co-culture or mono-culture grown in the indicated conditions: 10x Ker: normal co-culture conditions with 10x keratinocytes in normal co-culture growth media; Mel-MM: melanocytes in mono-culture grown in melanocyte growth media; Mel-KCM: melanocytes in mono-culture grown in keratinocyte conditioned co-culture media. (**D**), Percent of GCaMP6f-expressing melanocytes with at least one Ca^2+^ transient grown in the indicated conditions. One way ANOVA F(2,9)=59.68, p-value <0.00001; post hoc Tukey means comparison to other culture conditions, ** p-value <0.00005; ns, not significant: p-value > 0.05. Cultures were derived from four skin donors as indicated by the different colored. All scale bars 200 µm.

**Figure 4 supplement 2:**
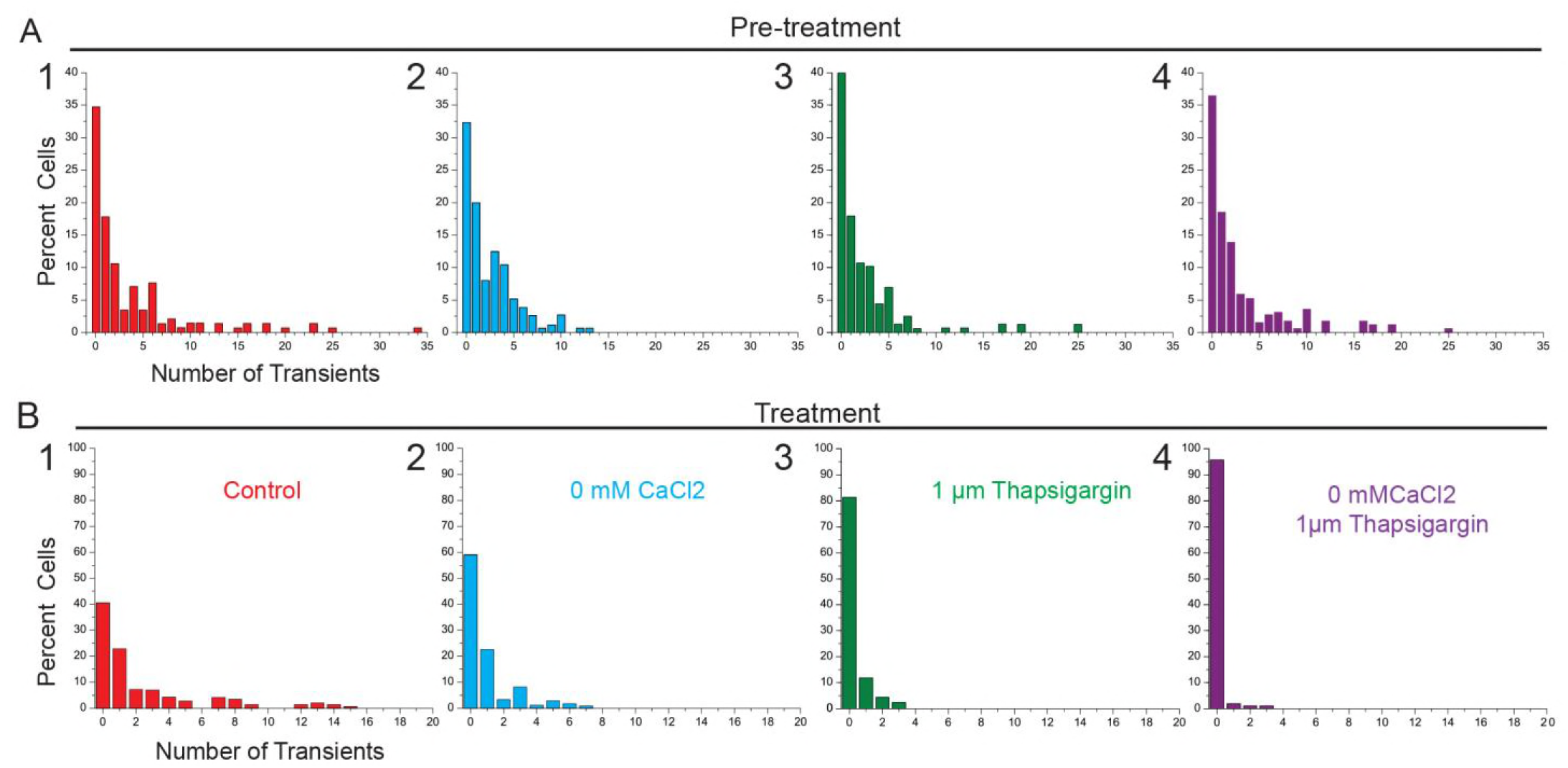
Distribution of the number of Ca^2+^ transients per cell changes upon removal of external and/or internal Ca^2+^ stores. (**A**), Normalized percent of cells with indicated number of Ca^2+^ transients from co-cultures derived from 3 skin donors in Figure 4. (**B**), Data from baseline imaging before treatment. **i**, Distribution of the number of Ca^2+^ transients per cell after the following treatments: 1) control with no treatment 2) CaCl_2_ removal 3) addition of 1 µM thapsigargin and 4) CaCl_2_ removal with addition of 1 µM thapsigargin. Colors in (**B**) correspond to conditions labeled in (**A**).

**Figure 4 supplement 3:**
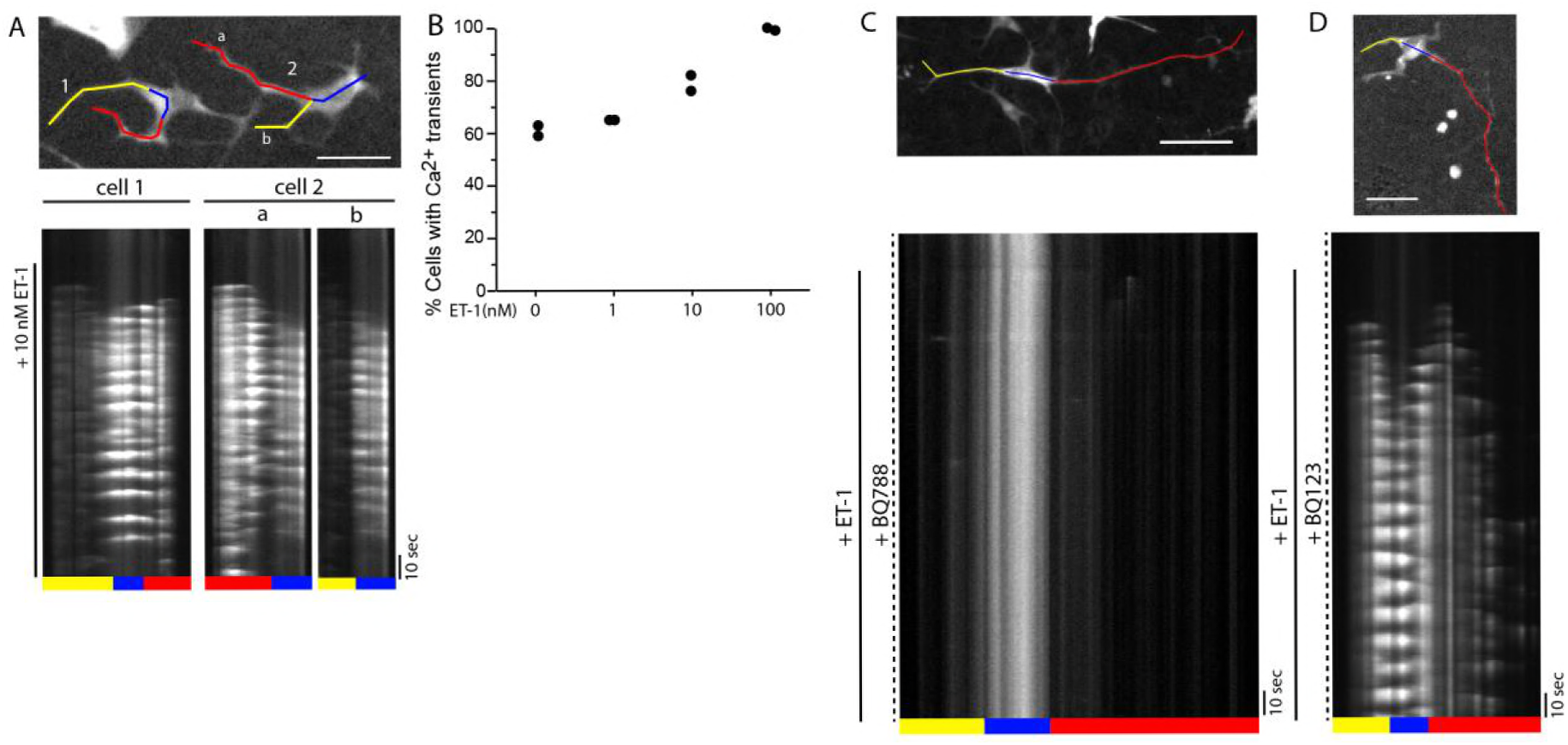
ET_B_ mediated dose dependent response to ET-1 in melanocytes co-cultured with keratinocytes (**A**), 10nM ET-1 induces Ca2+ transients across the entire melanocyte. Top panel: 2 representative melanocytes (cell 2 is the same cell from Figure 4C). Scale bar 100 µm. Bottom panel: Kymographs from corresponding color coded lines in top panel; black line denotes change to imaging media containing 10nM ET-1. (**B**), Percent of melanocytes with Ca2+ transients in the presence of 0, 1, 10 or 100nM ET-1 during a 2.5 imaging. N= 2 co-cultures from 2 different donor skin cells. (**C**), ET_B_ antagonist BQ788 abrogates ET-1 induced Ca2+ response in co-cultured melanocytes. Top panel: representative melanocyte Scale bar 150 µm. Bottom panel: Kymograph from corresponding color coded line in top panel; solid black line denotes change to imaging media containing 10nM ET-1 while maintaining 1 µM BQ788 (dashed black line). (**D**), BQ123 inhibition of ET_A_ did not inhibit melanocyte Ca2+ response to 10nM ET-1. Top panel: representative melanocyte, scale bar 100 µm. Bottom panel: Kymograph from corresponding color coded line in top panel; solid black line denotes change to imaging media containing 10nM ET-1 while maintaining 1 µM BQ123 (dashed black line).

**Figure 4 supplement 4:**
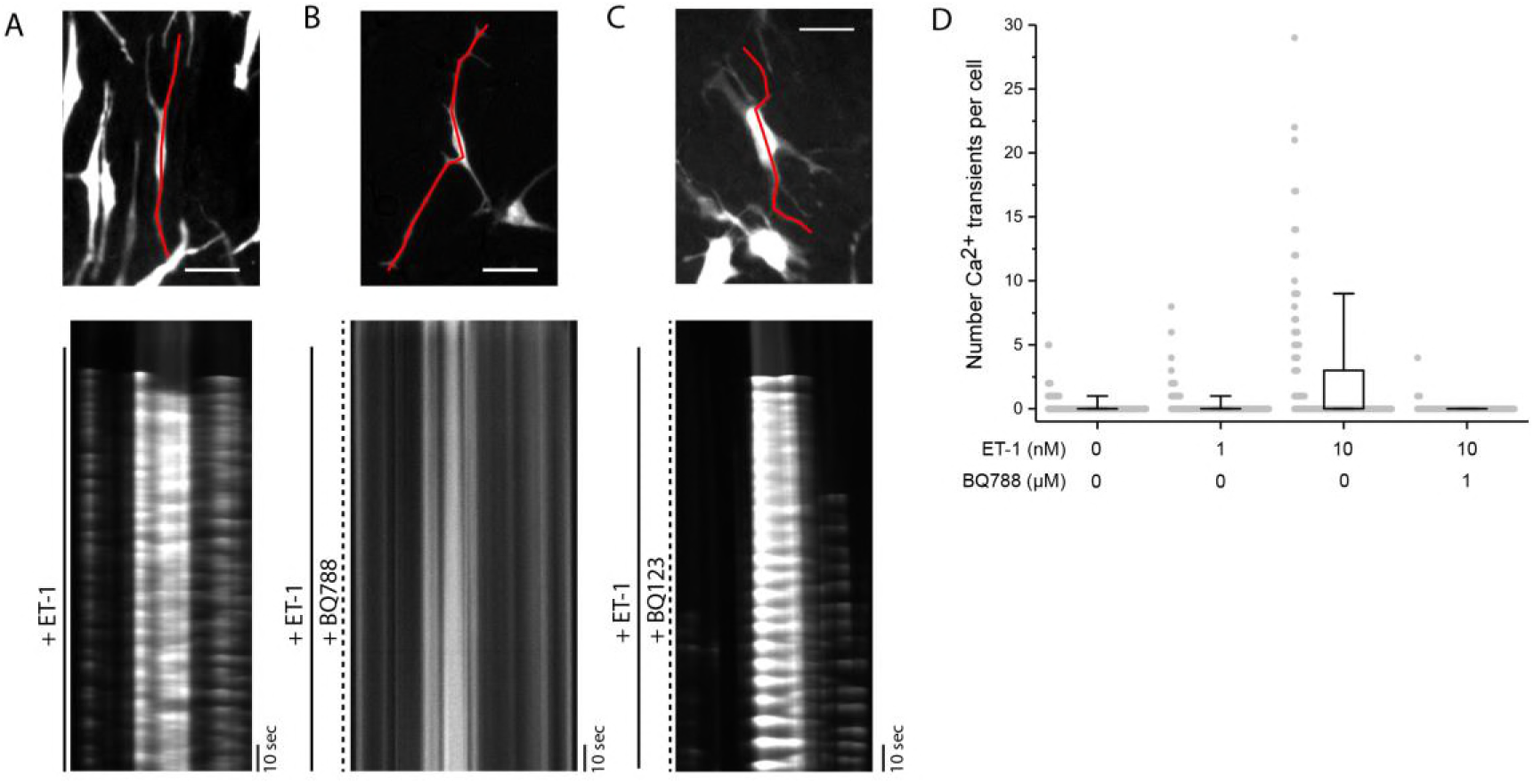
ET_B_ mediated dose dependent response to ET-1 in melanocytes mono-culture. (**A**), 10nM ET-1 induces Ca2+ oscillations across the entire melanocyte. Top panel: representative melanocyte. Bottom panel: Kymographs from corresponding line in top panel; black line denotes change to imaging media containing 10nM ET-1. (**B**), ET_B_ antagonist BQ788 abrogates ET-1 induced Ca2+ response in mono-cultured melanocytes. Top panel: representative melanocyte. Bottom panel: Kymograph from line in top panel; solid black line denotes change to imaging media containing 10nM ET-1 while maintaining 1 µM BQ788 (dashed black line). (**C**), BQ123 inhibition of ET_A_ did not inhibit melanocyte Ca2+ response to 10nM ET-1. Top panel: representative melanocyte Bottom panel: Kymograph from corresponding line in top panel; solid black line denotes change to imaging media containing 10nM ET-1 while maintaining 1 µM BQ123 (dashed black line). (**D**), Number of Ca2+ transients per mono-cultured melanocyte during 2.5 minutes in the presence of 0nM ET-1, 1nM ET-1, 10nM ET-1 or 10nM BQ788 (n=159,107,107,52 cells from 2, 1, 1, and 1 cultures respectively). All scale bars 100 µm.

**Figure 4 supplement 5:**
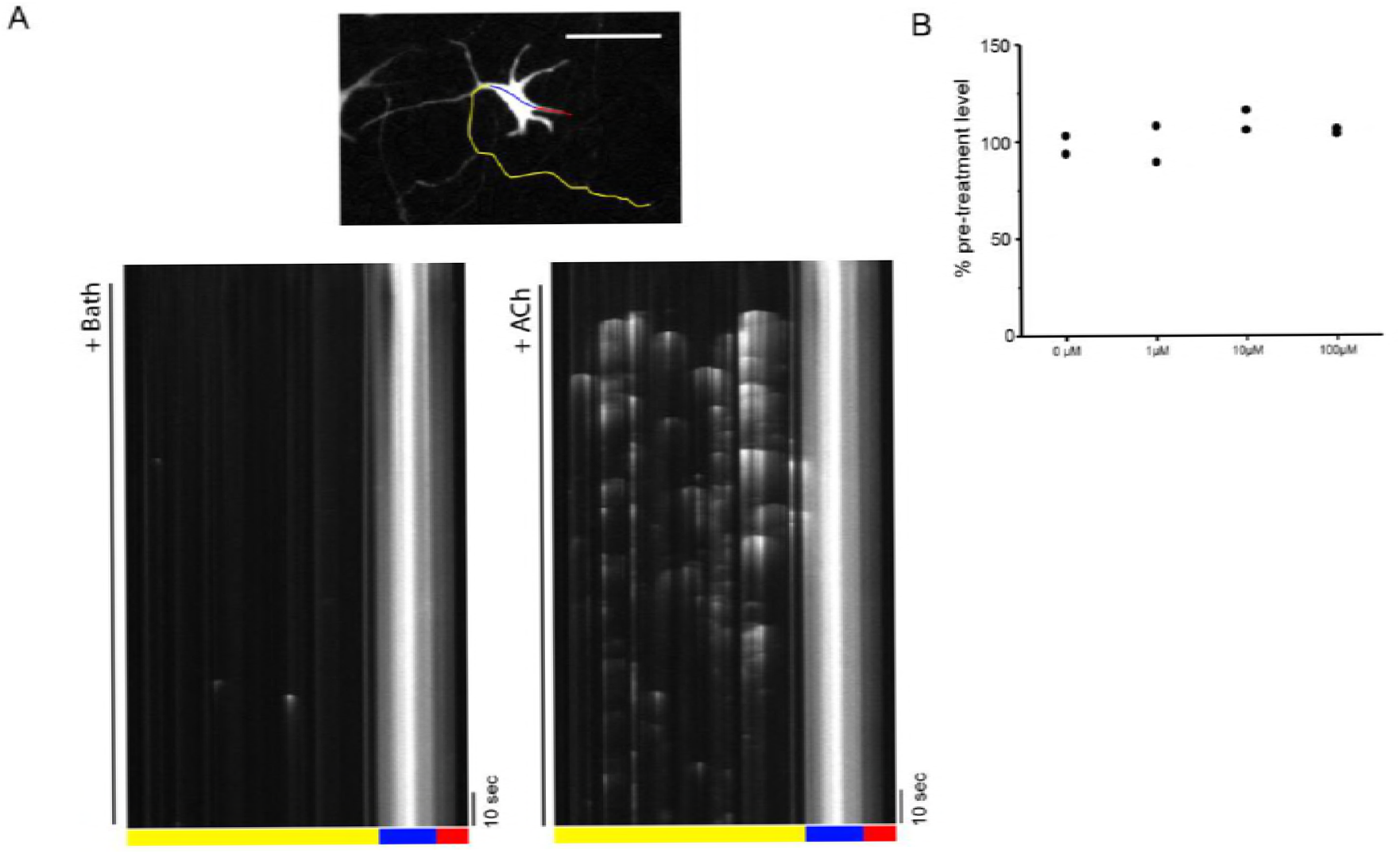
Melanocyte response to ACh in co-culture. (**A**), Representative melanocyte, in co-culture with keratinocytes, with Ca^2+^ response to 100 µM ACh. Bottom panels: Kymograph from corresponding color coded line in top panel; black line denotes change to fresh imaging media (left, “bath”) or change to imaging media containing 100 µM ACh (right, +ACh). (**B**), ACh does not increase the number of melanocytes with Ca^2+^ transients during 2.5 minutes, n = 2 co-cultures from different donor skin cells per condition.

**Figure 4 supplement 6:**
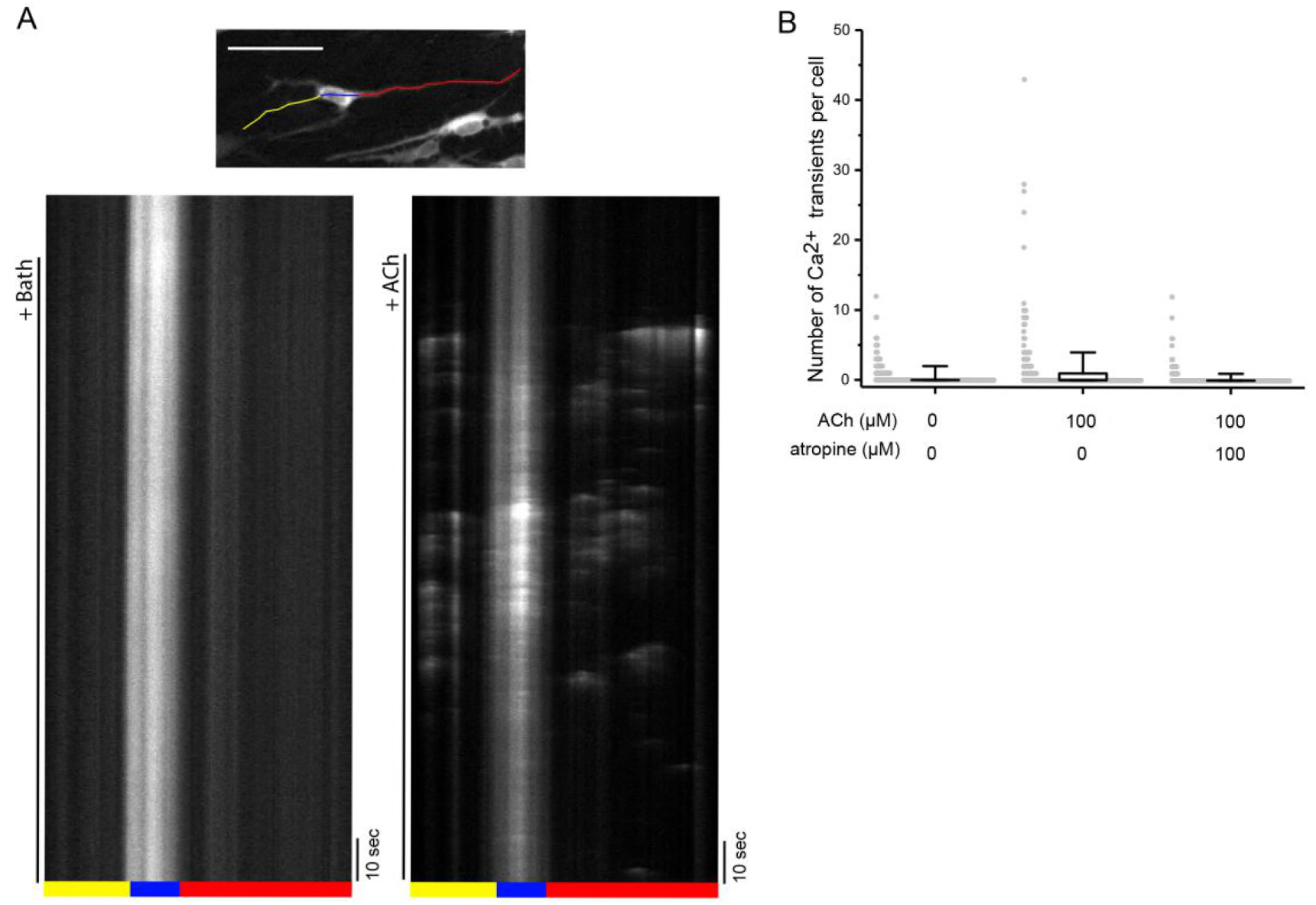
**Mono-cultured** melanocyte response to ACh. (**A**), Representative melanocyte, from mono-culture, with Ca2+ response to 100 µM ACh. Bottom panels: Kymograph from corresponding color coded line in top panel; black line denotes change to fresh imaging media (left, “bath”) or change to imaging media containing 100 µM ACh (right, +ACh). Scales bar 150 µm. (**B**),Number of Ca2+ transients per mono-cultured melanocyte during 2.5 minutes in the presence of 0 µM ET-1, 100 µM ET-1, or 100 µM atropine (n=405, 203, 202 cells from 4, 2, and 2 cultures from different donor skin cells, respectively). All scale bars 150 µm.

Movie 1: Ca2+ Transients in melanocytes co-cultured with keratinocytes. Related to Figure 3.

Movie 2: Ca2+ transients in co-cultured melanocyte dendrite and dendritic spine. Related to Figure 3C.

Movie 3: Ca2+ transient in co-cultured melanocyte dendritic spines. Related to Figure 3. Scale bar: 2µm..

Movie 4: Co-cultured melanocyte dendritic Ca2+ transients. Related to figure Figure 3E,F.

Movie 5: Melanocyte response to 10nM ET-1 in co-culture with keratinocytes. Related to Figure 4C..

Movie 6: BQ788 but not BQ123 inhibits co-cultured melanocyte Ca2+ response to ET-1Movie 7: Co-cultured melanocyte response to 100 µM Ach. Related to Figure 4E, Figure 4 Supplement 3

Movie 7: Co-cultured melanocyte response to 100 µM Ach. Related to Figure 4H, Figure Supplement 5.

## Materials and Methods

### Primary Cell Isolation

Melanocytes and keratinocytes were isolated as previously described ^31^ from neonatal foreskins obtained after routine circumcision, provided by Mount Sinai School of Medicine, New York, NY or Weill Cornell Medical College, New York, NY, in accordance with the Rockefeller University IRB protocol RBE-0721. Tissue was kept in CO_2_ Independent Media (Gibco - ThermoFisher Scientific, Carlsbad, CA) with 1x Antibiototic-Antimycotic (Gibco) at 4˚ C until processed for cell culture. In short, the epidermis was removed from the dermis via 10 mg/ml dispase II, neutral protease, grade II (Roche – Sigma, St. Louis, MO) digestion for 14 -17 hours at 4˚ C. Single cell suspension was achieved by mincing the epidermis and digesting with 0.5% trypsin (Gibco) for 5 minutes at 37˚ C. Trypsin was deactivated with soy bean trypsin inhibitor (Gibco) and the cells were washed with Hanks' Balanced Salt Solution, no Mg^2+^, no Ca^2+^ (Gibco) before plating. Cells were grown at 37˚C, in 5% CO_2_ with 1x Antibioltic-Antimycotic. After the first media replacement all antibiotics and antifungals were omitted from the growth media.

### Cell Culture

Melanocytes and keratinocytes were grown in melanocyte growth media (Medium 254 with Human Melanocyte Grow Serum-2 (HMGS-2), Gibco) and keratinocyte growth media (Epilife with Human Keratinocyte Growth Serum (HKGS), Gibco), respectively. Only cells 5 passages or less were used for experiments. 293T-HEK cells were grown in DMEM, high glucose, pyruvate media (Gibco) supplemented with non-essential amino acids (Gibco) and 10% FBS at 37˚C, 5% CO_2_.

#### Optimized Melanocyte – Keratinocyte Co-culture System

Building upon previous studies (33-35), we optimized a melanocyte-keratinocyte co-culture to mimic physiological conditions that also allowed us to generate the culture within a few days and use standard wide-field fluorescence microscopy techniques.

A confluent multi-layer sheet of melanocytes and keratinocytes was achieved by seeding at a high density (0.5× 10^4^ melanocytes and 5×10^4^ keratinocytes per 78 mm^2^) and allowing the cells to adhere and grow at 37˚C, 5% CO_2_ for 17-24 hours in keratinocyte growth media (EpiLife media with HKGS). This media formulation is used for mono-culture of primary human keratinocytes without the presence of a fibroblast feeder layer and contains 0.06mM CaCl_2_. This lower than physiological CaCl_2_ is used to maintain keratinocytes in an undifferentiated state which allows them to proliferate^32,33^. Melanocytes will grow and survive in keratinocyte media, in the presence of keratinocytes. However, both cell-cell adhesion and keratinocyte differentiation is Ca^2+^ dependent ^34^. Co-cultures kept at low CaCl_2_ remained as a monolayer, did not form adherens junctions and exhibited an altered cytoskeletal morphology that differed from healthy human skin (Figure 2 Supplement 1A). To achieve a multi-layer culture with proper cell-cell adhesion, cell morphology and both undifferentiated and differentiating keratinocytes, we increased the CaCl_2_ to 1.06 mM in the growth media after the initial monolayer was formed. This restored adherens junctions and supported a more physiological actin cytoskeleton structure in the keratinocytes (Figure 2 Supplement 1A,B). Within 48 hours after media replacement, co-cultures contained two populations of keratinocytes: undifferentiated K14 positive keratinocytes and larger K10 positive keratinocytes (Figure 2 Supplement 1C). Consistent with what is seen *in vivo,* the K14 positive keratinocytes were located at the basal surface of the culture while the K10 positive keratinocyte were located above the K14 positive cells.

Melanocytes in the co-cultures grown in 1.06 mM CaCl_2_ keratinocyte growth media, exhibited morphologies similar to those of melanocytes *in-situ* (Figure 2, Figure 2 Supplement 1D).

In short: For melanocyte – keratinocyte co-cultures, melanocytes and keratinocytes were seeded at a 1:10 ratio (0.5×10^4^ : 5×10^4^ cells) onto the 10 mm glass window of a 35 mm MatTek dish (P35G-1.5-10-C, MatTek, Ashland, MA) in keratinocyte growth media and incubated 12-24 hours at 37˚ C, 5% CO_2_. Once a confluent monolayer was achieved the media was changed to keratinocyte growth media containing 1.06 mM CaCl_2_ final concentration. Subsequent experiments were performed 48-72 hours after switching to 1.06mM CaCl_2_ media.

### Immunofluorescence

#### Frozen Sections

Within 12 hours after harvest, neonatal foreskin (kept at 4˚C in DMEM, high glucose, no pyruvate (Gibco) and Antibiotic-Antimytotic or CO_2_ Independent media (Gibco) and Antibiotic-Antimytotic) was fixed overnight at 4°C with 4% paraformaldehyde (Electron Microscopy Sciences, Hatfield, PA). After washing with PBS, the fixed tissue was incubated with 30% sucrose in DPBS over night at 4˚C. The sucrose was exchanged with OCT (VWR, Radnor, PA) and frozen on dry ice. In Figure S1B, a cryostat section of approximately 5-10 µm was used. After immunofluorescence, the section was mounted in ProLong^®^ Gold (Molecular Probes) and sealed.

#### Epidermal Sheets

For Figures 1A and Figure 1 Supplement 1, the epidermis was removed from the dermis by overnight incubation with dispase II. Freshly isolated epidermal sheets were fixed with 4% paraformaldehyde over night at 4˚C and washed before immunofluorescence staining. Epidermal sheets were imaged in PBS rather than mounting media to prevent compaction of tissue and loss of z resolution.

#### Cell Cultures

Cell cultures were wash 1X with PBS, fixed in 4% paraformaldehyde for 10 – 30 minutes at room temperature, and then washed before immunofluorescence staining.

After fixation, tissue (either fixed frozen sections or epidermal sheet) or cells were incubated in blocking buffer: 2.5% Donkey Serum, 2.5% Goat Serum (Jackson ImmunoResearch Laboratories, West Grove, PA), 1% Bovine serum albumin (Sigma), 0.1% Triton X-100 (Sigma) for 1 – 2 hours at room temperature. The following primary antibodies where used at the indicated concentration in blocking buffer overnight at 4˚C : anti-TRP1 (TA99, ab3312) 1:200, anti-K14 (LL002, ab7800) 1:300, anti-E cadherin (ab15148) 1:100 (Abcam, Cambridge, UK), anti-cKit 1:100 (CD11705, Invitrogen-Thermo Fischer Scientific), anti-K10 1:500 (Poly19054, BioLegend, San Diego, CA). Secondary antibodies against mouse IgG or rabbit IgG conjugated to Alexa Fluor 488, 594, or 680 (Molecular Probes – Thermo Fischer Scientific) were used at a 1:1000 dilution for 1 – 2 hours at room temperature. Hoechst (Molecular Probes) was used at 1:1000 dilution in PBS. For two color double anti-TRP1 and anti-cKit immunofluorescence (Figure S1A), two sequential rounds of primary and secondary antibodies were used with a 1.5 hours non-labeled secondary anti-mouse IgG (Molecular Probes) incubation in-between to insure the 2^nd^ secondary antibody did not have access the first primary antibody.

### Microinjections

Freshly isolated neonatal foreskin epidermis was obtained by an overnight dispase digestion at 4˚C and removal from the dermis while one ice. Individual melanocyte were microinjected using a manual microinjection (model MM-88-0, Narashige International Inc, East Meadow, NY) system on an Upright BX61WI microscope with manual syringe pressure application. 2% (w/v) Lucifer Yellow CH, Lithium Salt (Molecular Probes) was directly injected into the basal surface of the melanocyte at room temperature.

### Electron Microscopy

#### FIB-SEM

Neonatal foreskin was immersion fixed in 4% paraformaldehyde, 2% glutaraldehyde, 2mM CaCl_2_, 0.1M sodium cacodylate at 4°C. Tissue was cut into 1 mm x 500 µm pieces for processing. Tissue was then post fixed with 2% osmium tetroxide, 1.5% potassium ferrocyanide on ice for 1 hour followed by a thiocarbohydrazide amplification step for 20 minutes and then a subsequent 2% osmium tetroxide incubation for 30 minutes at room temperature. Tissue was then stained with 1% uranyl acetate overnight at 4°C. Lead aspartate staining was performed for 30 minutes at 60°C the next day followed by tissue dehydration using sequential ethanol dehydration followed by propylene oxide and embedded in Durcapan resin (Sigma-Aldrich). The sample block was trimmed then mounted on a SEM sample holder using double-sided carbon tape (EMS). Colloidal silver paint (EMS) was used to electrically ground the exposed edges of the tissue block. The entire surface of the specimen was then sputter coated with a thin layer of gold/palladium. The tissue was imaged using back scattered electron (BSE) mode in a FEI Helios Nanolab 650. Images were recorded after each round of ion beam milling using the SEM beam at 2 keV and 100 pA with a working distance of 2.8 mm. Data acquisition occurred in an automated way using the Auto Slice and View G3 software. The raw images were 2,048 1,768 px2, with 30 nm slices viewed at a -38 degree cross-sectional angle. Each raw image had a horizontal field width (HFW) of 30 µm with an XY pixel size of 14-18 nm and 30 nm Z slices. These images were then aligned using the image processing programs in IMOD ^35^. The aligned images were then use to generate a 3-D model in IMOD’s 3dmod image display and modeling program.

#### TEM serial sections

Neonatal foreskin was immersion fixed in 4% paraformaldehyde, 2% glutaraldehyde, 2mM CaCl_2_, 0.1M sodium cacodylate at 4°C. Tissue was cut into 1 mm x 500 µm pieces for processing. Tissue was post fixed with 1% osmium tetroxide on ice for 3 hours followed by uranyl acetate staining (1% in 0.05 M maleate buffer pH 5.2) overnight at 4°C. The next day, tissue was dehydrated using sequential ethanol dehydration followed by propylene oxide and embedded in Epon resin (Electron Microscopy Sciences). “Grey sections” for TEM (approximately 50 -100 nm sections) were used. Sections were imaged on a JEOL JEM 100CX Transmission Electron Microscope (JEOL USA, Peabody, MA) with AMT 2K x 2K digital camera (Woburn, MA) at the Rockefeller University Electron Microscopy Resource Center.

### Keratinocyte – Melanocyte contact dependence assay

To assess the percent of melanocytes with transients in co-culture and mono-culture conditions, all cultures were seeded onto the 10 mm glass window of a collagen I coated MatTek dish. The follow seeding ratios and growth media were used: “10x ker” is 0.5×10^4^ melanocytes expressing GCaMP6f (mel6f) and 5×10^4^ keratinocytes, grown in keratinocyte growth media; “Mel alone” is 0.5×10^4^ mel6f, grown in melanocyte media; “10x Mel” is 0.5×10^4^ mel6f and 5×10^4^ non-transduced melanocytes in melanocyte media; “10x 293T” is 0.5×10^4^ mel6f and 5×10^4^ HEK293T cells; “TW” is 0.5×10^4^ mel6f seed on glass bottom of Collagen I coated MatTek dish with 5×10^4^ keratinocytes seed into a collagen I coated Transwell culture insert, 3 µm pore (CORNING, Corning, NY) above the glass window, grown in keratinocyte media. For all culture conditions, cultures were incubated over night at 37˚ C, 5% CO_2_ and media was replaced with fresh media containing 1.06 mM CaCl_2_. Cultures were imaged 48 hours after media change. To compare the effect of different growth conditions for co-cultures and mono-cultures, 10x Ker, Mel alone, 10x Mel, 10x 293T, TW 10x Ker were grown in triplicate with one dish for each media condition: keratinocyte media, melanocyte media and 50/50 media (1:1 melanocyte:keratinocyte media). Keratinocyte conditioned media was obtained by harvesting the supernatant from a skin donor matched keratinocyte culture at each step of the culturing process mentioned above.

### Image Acquisition and Quantification

#### Immunofluorescence

Images were acquired using cellSens software (Olympus, Waltham, Massachusetts) on an IX83 microscope (Olympus) with a UPLSAPO 60× 1.3 NA silicone oil objective (Olympus), and a Orca-ER CCD digital camera (Hamamatsu Photonics, Bridgewater, NJ). Deconvolution was done with Autoquant (Media Cybernetics, Rockville, MD) standard settings for each filter set using blind adaptive point-spread function. Image analysis was done in Fiji (http://fiji.sc/).

#### Cell cultures

Cultures in MatTek dishes were imaged (fluorescence and IR-DIC) on an Upright BX61WI microscope (Olympus) with a UMPlan FL 60x 1.0 NA water dipping objective(Olympus), Orca Flash 4.0 digital CMOS camera (Hamamatsu) using MetaMorph image acquisition software (Molecular Devices) or a DeltaVision system (GE Healthcare, Pittsburgh, PA) at the Rockefeller University Bio-Imaging Resource Center with a 60x N.A. 1.52 oil objective and SoftWoRx software (DeltaVision). Deconvolution was performed using the adaptive point spread setting in Autoquant (Media Cybernetics) or, for images acquired on the DeltaVision, deconvolution was performed using the measured point-spread function in the SoftWoRx software. For live cell Imaging, cultures were kept in co-culture growth media at room temperature. Image analysis was done in Fiji.

#### Ca^2+^ Imaging

Prior to imaging, melanocyte – keratinocyte co-cultures were washed 3x with modified DPBS (10 mM Hepes, 10 mM D-Glucose, 0.1µM Glycine, 0.5 mM MgCl_2_, pH 7.1-7.3) with 1.06 mM CaCl_2_ added. Cultures were imaged in streaming acquisition mode in Metamorph Software (Molecular Devices) with a 250 ms exposure time on the Upright BX61WI microscope at room temperature using UMPlan FL 10x, 0.3 NA water dipping objective or UMPlan FL 60x, 1.0 NA water dipping objective.

*Number of cells with transients* was quantified by manually counting the cells with at least one transient Ca^2+^ signal during the 2.5 minute imaging period. For each positive cell, presence of local or global Ca^2+^ transients was manually determined.

*Number of transients per cell* were quantified by manually counting individual peaks in fluorescence by eye using line scans along the dendrite or an ROI was drawn at initiation sight and a plot of the fluorescence intensity over time was used to determine the number of peaks and thus the number of transients.

*Spatiotemporal assessment of transients* was done on images acquired in streaming mode with 50 ms or 250 ms exposure using a UMPlan FL 60x, 1.0 NA water dipping objective. The distribution of the spread in Ca^2+^ signal was determined using data acquired in streaming mode with a 250 ms exposure time using a UMPlan FL 10x, 0.3 NA water dipping objective.

*Calculating ∆F/F0:* F0 was determined by averaging 10 consecutive fluorescence intensities, within the same ROI, at the lowest fluorescence intensity point during the imaging sequence. ∆F was determining by: ∆F = Fn – F0, where Fn is the fluorescence intensity at that given time point. For ∆F/F0 images: 10 consecutive frames were averaged and the resulting image was used as F0. The ∆F image was obtained by subtracting the F0 image from the frame corresponding to the peak Ca^2+^ signal in corresponding ∆F/F0 plot. The final ∆F/F0 image was obtained by dividing the ∆F image by the F0 image.

### Ca^2+^ Source, Agonist and Antagonist Assays

Co-cultures in modified DPBS, 1.06 mM CaCl_2_, were imaged for 2.5 minutes in streaming mode, 250 ms exposure, to determine the baseline percentage of cells with transients and the number of transients per cell. After baseline imaging, media was replaced with modified DPBS, containing no added CaCl_2_ and/or 1 µM thapsigargin, 1 µM BQ788, 1 µM BQ123 (Tocris – BioTechne, Minneapolis, MN), 100 µM atropine sulfate (Sigma) or 100 µM atropine sulfate with 1 µM BQ788. To insure full replacement of the imaging media with the treatment solution, a 10x excess of the solution was perfused into the imaging chamber of a MatTek dish using manual syringe driven delivery through a modified open perfusion insert (model #RC-37F, Warner Instruments, Hamden, CT). The same field of view imaged for the baseline analysis was imaged after a 20-minute incubation with the perfused solution. The fold change in percent melanocytes with one or more Ca^2+^ transient was determined by dividing the number of cells with transients during the treatment imaging period by the number of cells with transients during the baseline imaging period. For agonist stimulation, acetylcholine chloride (Tocris) or endothelin-1 acetate salt (Bachem) was perfused into the chamber at 10x excess volume, to insure complete replacement of imaging media, at the indicated time during the streaming acquisition.

### Lentiviral vectors and production

*Phi3-EGFPmem:* EGFPmem was made by replacing ECFP with EGFP in the Clontech vector pECFPmem (Clontech – Takara Bio USA, Mountain View, CA). EGFPmem was inserted into the HIV-1 based lentivirus vector pLVX Phi3 ^36^ using restriction enzymes BamHI and XbaI and Quick Ligation Kit (New England BioLabs, Ipswich, MA).

*Phi3-iRFPmem:* Annealed sense and antisense DNA oligos containing the sequence for the first 20 amino acids of neuromodulin (mem) flanked by 5’ XhoI and 3’ NotI restriction enzyme sites were inserted into pLVX Phi3 by restriction digest and subsequent ligation using the Quick Ligation Kit (New England BioLabs). iRFP ^37^ was PCR amplified from piRFP (a gift from Vladislav Verkhusha) using Forward primer: 5’ CAAAAGATCGCGGCCGAAGGATCTGTCGCCAGG 3’, Reverse Primer: 5’ GCGTCCGGAGCGGCCTTACTCTTCCATCACGCCG 3’, and then inserted into the NotI digested Phi3-mem plasmid after the neuromodulin plasma membrane binding domain using the In-Fusion HD Cloning Kit (Clonetech).

*Zhi3-mCherry:* mCherry (from pmCherry-N1, Clonetech) was inserted in PLVX Zhi3 ^38^ using restriction enzymes MluI and BamHI and Quick Ligation Kit (New England Biolabs).

*Phi3-GCaMP6f*: GCaMP6f from pGP-CMV-GCaMP6f (a gift from Douglas Kim) was excised from Addgene plasmid # 40753 using restriction enzymes BglII and XbaI and inserted into BamHI and XbaI digested PLVX Phi3 ^36^ using the Quick Ligation Kit.

Low-passage HEK293T cells (60-80% confluent) were co-transfected using polyethyleneimine (Sigma) at a 5:5:1 ratio of a lentiviral plasmid, a HIV-1 GagPol plasmid, and a VSVg plasmid, respectively. To obtain viral stocks, 88 μg of total DNA was used to transfect a 150-mm dish of 70% confluent cells. Virus-producing HEK293Ts were maintained in DMEM, high glucose, pyruvate media (`Gibco) supplemented with non-essential amino acids (Gibco) and 3% FBS (Sigma). Virus-containing media was collected at 24 hours, 48 hours and 72 hours post transfection, filtered through a 0.45 μm filter (EMD Millipore, Billerica, MA) and concentrated with Lenti-X Concentrator (Clontech). Concentrated virus was re-suspended in Hanks' Balanced Salt Solution or phosphate buffered saline and stored in aliquots at 20x final concentration. High titer virus stocks were stored at -86˚C for future use.

### DsiRNA knock down

DsiRNAs were generated from DsiRNA sequences predesigned by IDT (Integrated DNA Technologies, Coralville, Iowa. Duplexed sequences were as follows: EDNRA 13.1 antisense: 5’ GGACAAGAACCGAUGUGAAUUACTT 3’ sense: 5’ AAGUAAUUCACAUCGGUUCUUGUCCAU 3’, EDNRA 13.2 antisense: 5’ GCAACCUUCUGCAUUCAUAAAUCT T 3’ sense: 5’ AAGAUUUAUGAAUGCAGAAGGUUGCUA 3’, EDNRB 13.1 antisense: 5’ CAUGUCAGUAUCAUGUUCUCUAAT T 3’ sense: 5’ AAUUAGAGAACAUGAUACUGACAUGGA 3’, EDNRB 13.2 antisense: 5’ AGUAUUGACAGAUAUCGAGCUGUT G3’ sense: 5’ CAACAGCUCGAUAUCUGUCAAUACUCA 3’. On day zero DsiRNA were introduced to melanocytes using Lipfectamine^®^ RNAiMAX transfection reagent (Invitrogen-Thermo Fischer Scientific), following the recommended transfection procedure for 35mm dishes. On day one, melanocytes were infected with Phi3-GCaMP6f and incubated overnight. On day two melanocytes were seeded with keratinocytes to generate co-cultures as described above.

### Statistical Analysis

Two sample T-test, One way ANOVA with Tukey means comparison and Kruskal-Wallis ANOVA statistical analyses were performed using standard settings in OriginPro software (OriginLab, Northampton, MA) as indicated in the figure legends and main text. We used the customary threshold of p < 0.05 to declare statistical significance. Sample size and statistical details can be found in the figures, legends and main text.

## Acknowledgements

We thank the Departments of Obstetrics and Gynecology at Mt. Sinai School of Medicine and Cornell-Weill for their assistance in the collection of the neonatal foreskins. We thank the Rockefeller Electron Microscopy Resource Center, The Rockefeller University Bio-Imaging Resource Center and the Simons Electron Microscopy Center for assistance. The FIB-SEM data was acquired at the Simons Electron Microscopy Center and National Resource for Automated Molecular Microscopy located at the New York Structural Biology Center, supported by grants from the Simons Foundation (349247), NYSTAR, and the NIH National Institute of General Medical Sciences (GM103310) with additional support from NIH S10 RR029300-01. We thank Pascal Maguin for assistance with cloning and Marina Bleck, Constantin Takacs, Michelle Itano, Sohail Tavazoie and Elaine Fuchs for helpful discussions. This work was supported in part by grant # UL1 TR001866 from the National Center for Advancing Translational Sciences (NCATS, National Institutes of Health (NIH) Clinical and Translational Science Award (CTSA) program, National Institute of General Medical Sciences of the National Institutes of Health under award number T32GM066699 and by a Rockefeller University Women in Science Fellowship to R.L.B.

## Author Contributions

R.L.B. performed the experiments and R.L.B. and S.S. designed the experiments and wrote the manuscript.

